# Adipocyte PI3K Links Adipostasis with Basal Insulin Secretion Through an Adipoincretin Effect

**DOI:** 10.1101/2023.04.24.538076

**Authors:** Barbara Becattini, Angela Molinaro, Marcus Henricsson, Jan Borén, Giovanni Solinas

**Affiliations:** Department of Molecular and Clinical Medicine, Institute of Medicine, University of Gothenburg, Gothenburg, Sweden; Department of Laboratory Medicine, Institute of Medicine, University of Gothenburg, Gothenburg, Sweden

## Abstract

Insulin secretion is governed by insulin-PI3K signaling. Resolving the mechanism of this feedback is necessary to understand how insulin operates. Mice lacking the insulin receptor, or AKT1 and AKT2 in adipocytes, are severely lipoatrophic. Thus, the role of adipocyte insulin-PI3K signaling in the control of insulin secretion remains unknown. Using adipocyte-specific PI3Kα knockout mice (PI3Kα^AdQ^) and a panel of isoform-selective PI3K inhibitors, we have found that PI3Kα and PI3Kβ activities are functionally redundant in adipocyte insulin signaling. PI3Kβ-selective inhibitors had no effect on adipocyte AKT phosphorylation in control mice but blunted AKT phosphorylation specifically in adipocytes of PI3Kα^AdQ^ mice, demonstrating adipocyte-selective inhibition of PI3K signaling. Adipocyte-selective PI3K inhibition increased serum FFA and potently induced insulin secretion. We name this phenomenon the adipoincretin effect. The adipoincretin effect was dissociated from blood glucose and blood glucose counterregulatory response. The contribution of lipolysis, lipid, and amino acid metabolism, and selected adipokines to the adipoincretin effect has been investigated. We conclude that basal insulin secretion is chiefly controlled by adipocyte PI3K signaling through the adipoincretin effect. This phenomenon reveals an essential role for adipocyte insulin-PI3K signaling in linking the rates of adipose tissue lipolysis with baseline insulin secretion during fasting.

## INTRODUCTION

Healthy humans with comparable blood glucose concentrations display a large variability in plasma insulin abundances (Kahn et al., 1993; Kodama et al., 2013), a phenomenon which is also observed between different strains of mice (Aston-Mourney et al., 2007). This capacity of different species to precisely match insulin secretion with insulin sensitivity over a wide range of serum insulin indicates a conserved crosstalk between the metabolic action of insulin and insulin secretion. The metabolic action of insulin depends on the induction of phosphoinositide-3 kinase (PI3K) - AKT signaling in insulin target cells (Solinas and Becattini, 2022). Pharmacological inhibition of either the insulin receptor (InsR), PI3K, or AKT causes hyperglycemia and hyperinsulinemia in humans and rodents (Ando et al., 2014; Hopkins et al., 2018; Kalinsky et al., 2018; Patnaik et al., 2016; von Mehren et al., 2020).

Thus, insulin secretion is chiefly controlled by a conserved negative feedback mediated by InsR-PI3K signaling in a cell type yet to be identified. This feedback is assumed to be mediated by insulin action on blood glucose, which involves hepatic glucose production (Vasan and Cantley, 2022). Indeed, the current paradigm for the insulin system can be regarded as glucose-centric, with blood glucose being considered the primary driver of insulin secretion.

However, whereas PI3K inhibitors induce a large and sustained increase in insulin secretion in fasted C57BL/6 mice (Hopkins et al., 2018), a glucose injection in these mice causes only a modest and transient increase in plasma insulin (Aston-Mourney et al., 2007; Fergusson et al., 2014; Small et al., 2022). Moreover, hepatocyte-restricted acute inhibition of PI3K activity, achieved by administering a PI3Kβ-selective inhibitor to mice lacking PI3Kα in hepatocytes, causes hyperglycemia without further increasing insulin secretion (Molinaro et al., 2019). Altogether, this evidence made us question the current glucose-centric paradigm for the insulin system and motivated us to search for an alternative mechanism for the feedback linking insulin action with insulin secretion.

In humans and rodents, β-adrenergic agonists stimulate lipolysis and insulin secretion independently from blood glucose (Imura et al., 1971; Porte, 1967; Sennitt et al., 1985), and inhibition of lipolysis with nicotinic acid (NA) reduces insulin secretion during fasting (Balasse and Ooms, 1973; Boden et al., 1998; Dobbins et al., 1998a; Dobbins et al., 1998b; Stein et al., 1996). Furthermore, we have found that in humans with different adiposity and insulin sensitivity, fasting plasma insulin was not associated with blood glucose but was associated with adipose tissue lipolysis (Fryk et al., 2021). Considering the opposing actions of β-adrenergic signaling and InsR-PI3K signaling on adipocyte lipolysis (Guilherme et al., 2022; Solinas and Becattini, 2022), we hypothesized that insulin secretion is governed by adipocyte InsR-PI3K signaling.

Mice lacking either the InsR or AKT1 and AKT2 in their adipocytes develop severe lipoatrophy and metabolic syndrome phenotype, demonstrating the requirement for functional insulin signaling in adipostasis (Boucher et al., 2016; Qiang et al., 2016; Sakaguchi et al., 2017; Shearin et al., 2016). However, because of their severely atrophic adipose tissue and metabolic syndrome phenotype, these models provide limited insights into the physiological role of InsR-PI3K signaling within the mature adipocyte. Thus, the role of adipocyte PI3K signaling in the feedback controlling insulin secretion remains unknown.

## RESULTS

### Adipocyte PI3Kα is dispensable for adipostasis and metabolic homeostasis in mice

It has been proposed that PI3Kα is the main PI3K catalytic isoform in adipocyte insulin signaling (Foukas et al., 2006; Knight et al., 2006). Therefore, we crossed PI3Kα LoxP floxed mice (PI3Kα^F/F^) (Molinaro et al., 2019) with mice expressing the Cre recombinase under the control of the AdipoQ promoter to generate mice lacking PI3Kα specifically in adipocytes (PI3Kα^AdQ^). Compared to PI3Kα^F/F^ mice, PI3Kα^AdQ^ mice showed a substantial reduction of PI3Kα protein abundance in white fat pads and a trend in interscapular brown fat, but not in other tissues (Figures S1A-D). Mature adipocytes purified from epididymal fat pads of PI3Kα^AdQ^ mice showed almost complete ablation of the PI3Kα protein without changes in the abundance of other catalytic or regulatory PI3K subunits or IRS1 and IRS2 (Figures S1E and F). PI3Kα protein abundance was not affected in adipose tissue stromal vascular cells of PI3Kα^AdQ^ mice (Figures S1G and H). Thus, PI3Kα protein was almost completely ablated in adipocytes of PI3Kα^AdQ^ mice but was comparable in abundance to control mice in adipose stromal vascular cells, muscle, liver, kidney, brain, and spleen (Figure S1).

PI3Kα^AdQ^ mice showed a normal growth curve, slightly reduced serum FFA levels, normal fasting glucose and insulin levels, normal glucose and insulin tolerance, and normal suppression of serum FFA levels by glucose (Figures 1A-C and S2A-C). Fat pad weights and adipocyte size distribution of PI3Kα^AdQ^ mice were similar to controls, and their liver histology was normal (Figures 1D-F and S2D). Therefore, in stark contrast with the phenotype of mice lacking InsR or AKT1 and AKT2 in their adipocytes (Boucher et al., 2016; Qiang et al., 2016; Sakaguchi et al., 2017; Shearin et al., 2016), PI3Kα^AdQ^ mice are not lipodystrophic but show normal adiposity, glucose, and lipid homeostasis. These results show that PI3Kα is functionally redundant with another PI3K isoform for insulin-PI3K-AKT signaling within the adipocyte.

**Figure 1.**
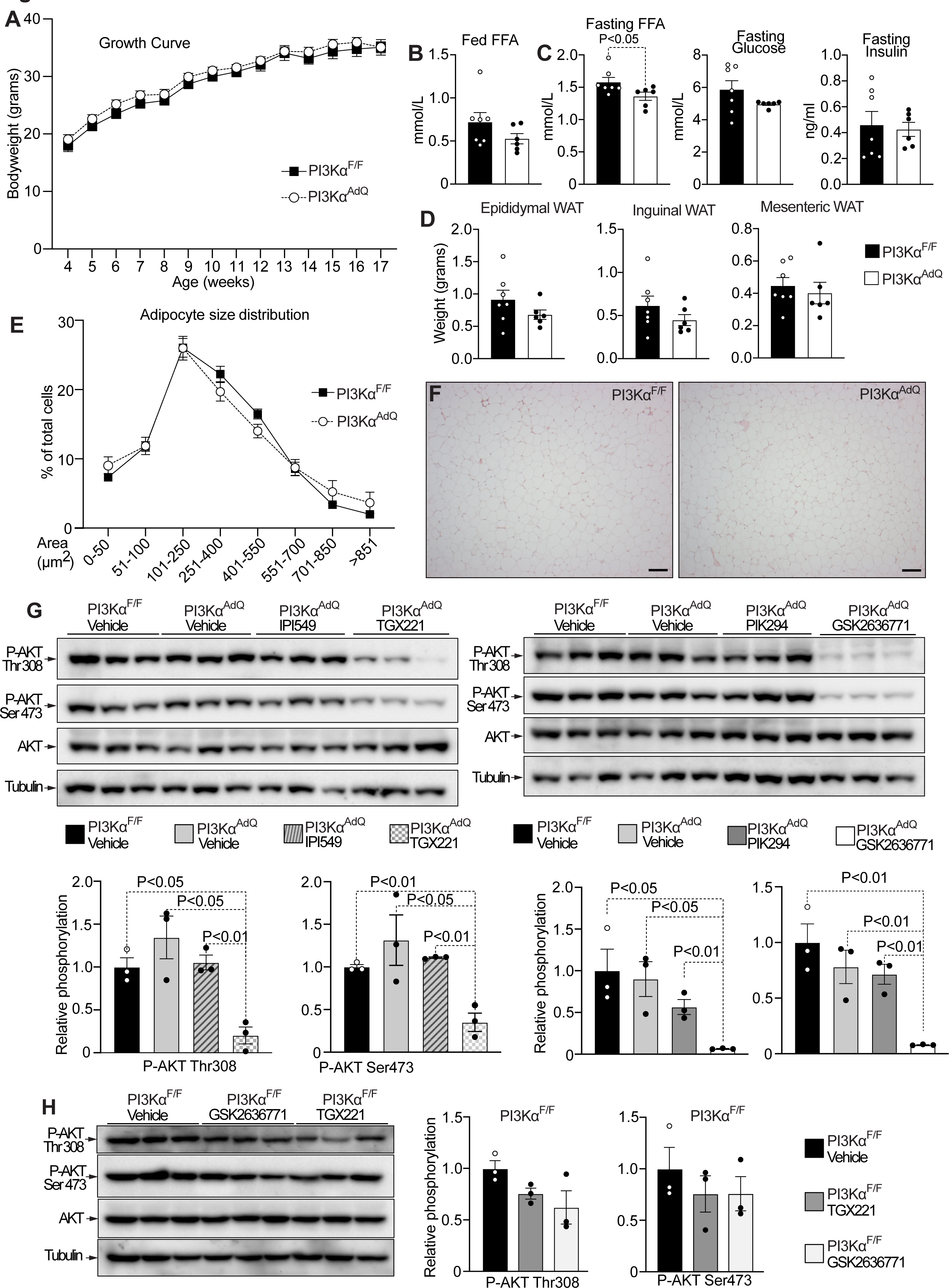
Adipocyte AKT signaling in-vivo depends on PI3Kα and PI3Kβ. (A) Growth curves of PI3Kα^AdQ^ and PI3Kα^F/F^ mice. (B) Fed serum free fatty acids (FFA) of the mice above. (C) Fasting serum FFA, fasting blood glucose, and fasting serum insulin of PI3Kα^AdQ^ and PI3Kα^F/F^ mice. (D) Adipose tissue weights of PI3Kα^AdQ^ and PI3Kα^F/F^ mice. (E-F) Adipocyte size distribution and representative histology of epidydimal fat pads. (G) In-vivo pharmacological mapping of the PI3K isoforms driving AKT signaling in the adipocyte of PI3Kα^AdQ^ and PI3Kα^F/F^ mice fasted for 12 hours and sacrificed 10 minutes after i.p. injection of 2 g/Kg of glucose with either vehicle or the indicated PI3K inhibitor: 6 mg/Kg for TGX221 (PI3Kβ), IPI549 (PI3Kγ), GSK2636771 (PI3Kβ), 12 mg/Kg for PIK294 (PI3K8). (H) The effect of the PI3Kβ-selective inhibitors TGX221 and GSK2636771 on AKT phosphorylation was evaluated in PI3Kα^F/F^ control mice. Scale bar 100 μm for F. n=7 mice PI3Kα^F/F^ and n=6 mice PI3Kα^AdQ^ for A-D, n=6 mice for E and F, 3 mice for G and H. Data are mean ± s.e.m. Statistical analysis was performed using Mann-Whitney for C or unpaired t-test for G.

### Adipocyte PI3K-AKT signaling depends on PI3Kα and PI3Kβ functional redundancy

To identify the PI3K isoforms driving AKT phosphorylation in the adipocyte, PI3Kα^F/F^ mice and PI3Kα^AdQ^ mice were fasted for 12 hours and received one intraperitoneal injection (i.p.) of 2 g/kg of glucose in the presence of different isoform-selective PI3K inhibitors or vehicle. Ten minutes after the injection, the mice were sacrificed, and the epididymal fat pads were collected to measure AKT phosphorylation. AKT phosphorylation was similar in the adipose tissues of PI3Kα^F/F^ mice and PI3Kα^AdQ^ mice injected with glucose and vehicle (Figure 1G). This demonstrates that, in adipocyte insulin-PI3K-AKT signaling, PI3Kα is functionally redundant with another PI3K isoform. The PI3Kγ-selective inhibitor IPI549 or the PI3K8- selective inhibitor PIK294 did not affect AKT phosphorylation in the adipose tissues of PI3Kα^AdQ^ mice. Thereby PI3Kγ and PI3Kδ activities are dispensable for insulin-PI3K-AKT signaling in adipocytes of PI3Kα^AdQ^ mice. However, two different PI3Kβ-selective inhibitors (TGX221 and GSK2636771) substantially reduced AKT phosphorylation in the adipose tissue of PI3Kα^AdQ^ mice, with GSK2636771 achieving more than 90% inhibition (Figure 1G). The PI3Kβ-selective inhibitors TGX221 and GSK2636771 had no significant effect on adipose tissue AKT phosphorylation in PI3Kα^F/F^ control mice (Figure 1H). Hence, AKT phosphorylation is not affected in cells with either a functional PI3Kα or PI3Kβ but it is blocked in adipocytes with inactivation of both PI3Kα and PI3Kβ.

These results demonstrate that insulin-PI3K-AKT signaling in the mouse adipocyte, in vivo, depends on functionally redundant PI3Kα and PI3Kβ activities.

### Adipocyte PI3K links adipostasis with insulin secretion

Our results are consistent with previous studies showing that PI3Kβ-selective inhibition does not interfere with insulin signaling (Foukas et al., 2006; Knight et al., 2006; Molinaro et al., 2019; Sopasakis et al., 2010). However, the results described above demonstrate that PI3K- AKT signaling can be inhibited acutely in adipocytes in vivo by administering a PI3Kβ- selective inhibitor to PI3Kα^AdQ^ mice (Figures 1G, H). An adipocyte-selective acute inhibition of PI3K signaling solves the problem of the lipoatrophy associated with genetic blockage of adipocyte InsR-PI3K-AKT signaling in mice (Boucher et al., 2016; Qiang et al., 2016; Sakaguchi et al., 2017; Shearin et al., 2016). Thus, we used this acute approach to investigate the role of adipose tissue PI3K signaling in the feedback control of insulin secretion. PI3Kα^F/F^ mice and PI3Kα^AdQ^ mice were fasted for 12 hours and received one intraperitoneal injection of 2 g/kg of glucose with either vehicle or the PI3Kβ-selective inhibitor GSK2636771. The experiment is acute and lasted 180 minutes from the injection, and the timepoint 0 minutes was just before the GSK2636771 injection. Blood glucose levels were similar between groups, and serum FFA levels were significantly reduced following the glucose injection in all the groups except for the PI3Kα^AdQ^ mice treated with the PI3Kβ- selective inhibitor GSK2636771, where serum FFA abundance was substantially increased (Figures 2A and B). Serum insulin and C-peptide (a marker for insulin secretion) were close to basal levels in all groups except for PI3Kα^AdQ^ mice treated with GSK2636771, where insulin secretion within 30 minutes was potently induced in a sustained manner (Fig. 2C and D). These findings reveal that low adipocyte PI3K activity triggers a feedback signal that potently induces insulin secretion. By analogy to the glucoincretin effect (Holst, 2019; Rehfeld, 2018), we named this phenomenon “the adipoincretin effect.” A similar, although less potent, effect was observed in PI3Kα^AdQ^ mice treated with the PI3Kβ-selective inhibitor TGX221 (Figures S3A-C). TGX221, which was less effective than GSK2636771 in reducing AKT phosphorylation in adipocytes of PI3Kα^AdQ^ mice (Figure 1G), blocked the action of glucose on FFA suppression but did not increase FFA above basal levels (Figure S3B).

**Figure 2.**
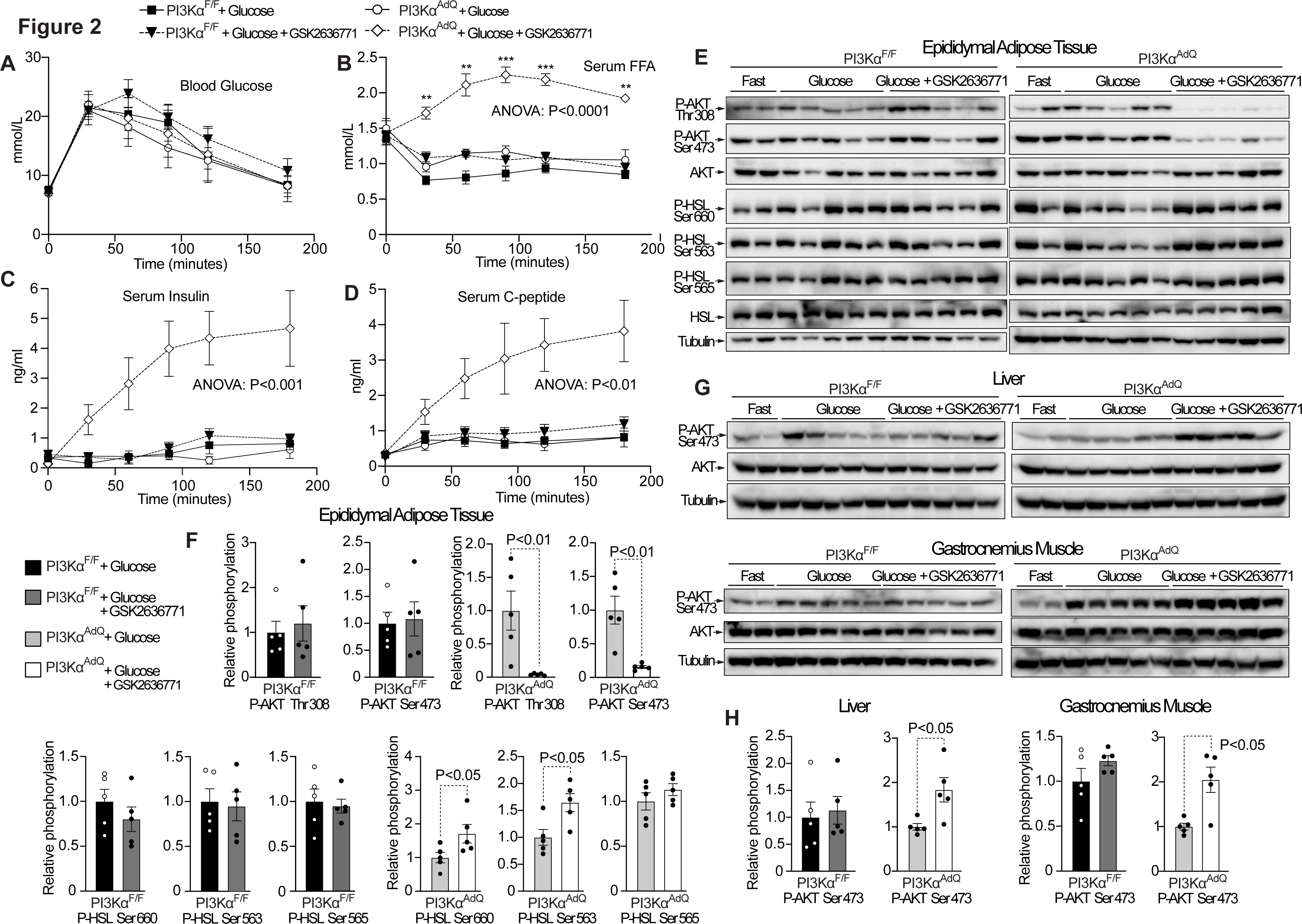
Adipocyte PI3K activity links adipostasis with insulin secretion through an adipoincretin effect. (A) Blood glucose was measured in PI3Kα^AdQ^ and PI3Kα^F/F^ mice fasted overnight for 12 hours and injected i.p. with 2 g/Kg of glucose with either vehicle or 6 mg/Kg of the PI3Kβ- selective inhibitor GSK2636771. (B) Serum FFA; (C) serum insulin; and (D) serum C-peptide of the mice from A. (E, F) immunoblot analysis of AKT and HSL phosphorylation in epididymal adipose tissue from mice as from A one-hour post-injection. (G, H) immunoblot analysis of AKT phosphorylation in liver and muscle from mice as from A one-hour post-injection. n=6 for A-D, n=2-5 for E, G (2 for fasting), n=5 for F, H. Data are represented as mean ± s.e.m. Statistical analysis was performed using repeated measures two-way ANOVA and Šídák’s multiple comparisons test for A-D, and Mann-Whitney for F,H. Indicated P-values in B-D are the most conservative for PI3Kα^AdQ^ + Glucose + GSK2636771 vs PI3Kα^AdQ^ +Glucose, or vs PI3Kα ^F/F^ + Glucose + GSK2636771.

Hence, the adipoincretin effect can partially be dissociated from increased serum FFA.

The PI3Kβ-selective inhibitors TGX221 and GSK2636771 did not affect insulin secretion in cultured islets from PI3Kα^F/F^ mice or PI3Kα^AdQ^ mice, which excludes a direct effect of PI3Kβ inhibition in islet cells on insulin secretion (Figure S3D).

To evaluate the efficacy of adipose tissue PI3K inhibition, a cohort of mice was sacrificed one hour after the GSK2636771 injection, and AKT phosphorylation was measured in different tissues. Since insulin-PI3K signaling reduces cAMP-PKA signaling in adipocytes (Solinas and Becattini, 2022), we have also measured the phosphorylation of hormone-sensitive lipase (HSL) at PKA and AMPK sites in adipose tissues of PI3Kα^AdQ^ mice and control mice. GSK2636771 showed no effect on AKT or ERK phosphorylation in PI3Kα^F/F^ mice but substantially reduced AKT phosphorylation, without affecting ERK phosphorylation, in the adipose tissues of PI3Kα^AdQ^ mice (Figures 2E, F, and S4). We observed increased AKT Ser-473 phosphorylation in the liver and muscle of PI3Kα^AdQ^ mice treated with GSK2636771 (Figures 2G and H), which is coherent with an adipose-specific PI3K inhibition and the increased insulin levels observed in these mice.

HSL phosphorylation at the PKA sites serine 660 and 563 was also significantly increased in the adipose tissues of PI3Kα^AdQ^ mice injected with GSK2636771 but not in control mice, whereas HSL phosphorylation at the AMPK site serine 565 was not altered (Figures 2E and F). In brown adipose tissue of PI3Kα^AdQ^ mice, GSK2636771 did not affect AKT phosphorylation but increased HSL phosphorylation at PKA sites (Figures S4E and F).

Overall, PI3Kβ-selective inhibition does not interfere with insulin signaling in cells expressing PI3Kα (e.g. muscle and liver), but selectively blocks AKT phosphorylation in adipocytes of PI3Kα^AdQ^ mice, where PI3Kα has been genetically ablated. This approach, which does not cause the severe lipoatrophy associated with genetic blockage of insulin- PI3K-AKT signaling in the adipocyte (Boucher et al., 2016; Qiang et al., 2016; Sakaguchi et al., 2017; Shearin et al., 2016), led us to discover that adipocyte PI3K signaling links insulin secretion with adipostasis through the adipoincretin effect.

### The adipoincretin effect is independent of glucose administration or feeding

Adipocyte-selective PI3K inhibition caused a potent and sustained insulin secretion, whereas an intraperitoneal injection of 2 g/kg of glucose in control mice did not cause an appreciable increase in serum insulin after 30 minutes from glucose administration (Figures 2A and C, D). This observation demonstrates that the adipoincretin effect cannot be fully explained by a rise in blood glucose. Therefore, we tested if the adipoincretin effect can be induced without administering the glucose injection in fed and fasting conditions.

Injection of GSK2636771, without glucose, significantly increased serum FFA and induced insulin secretion specifically in PI3Kα^AdQ^ mice but not in littermate control PI3Kα^F/F^ mice, in either fed conditions (Figures 3A-D) or after overnight fasting (Figures 3E-G).

**Figure 3.**
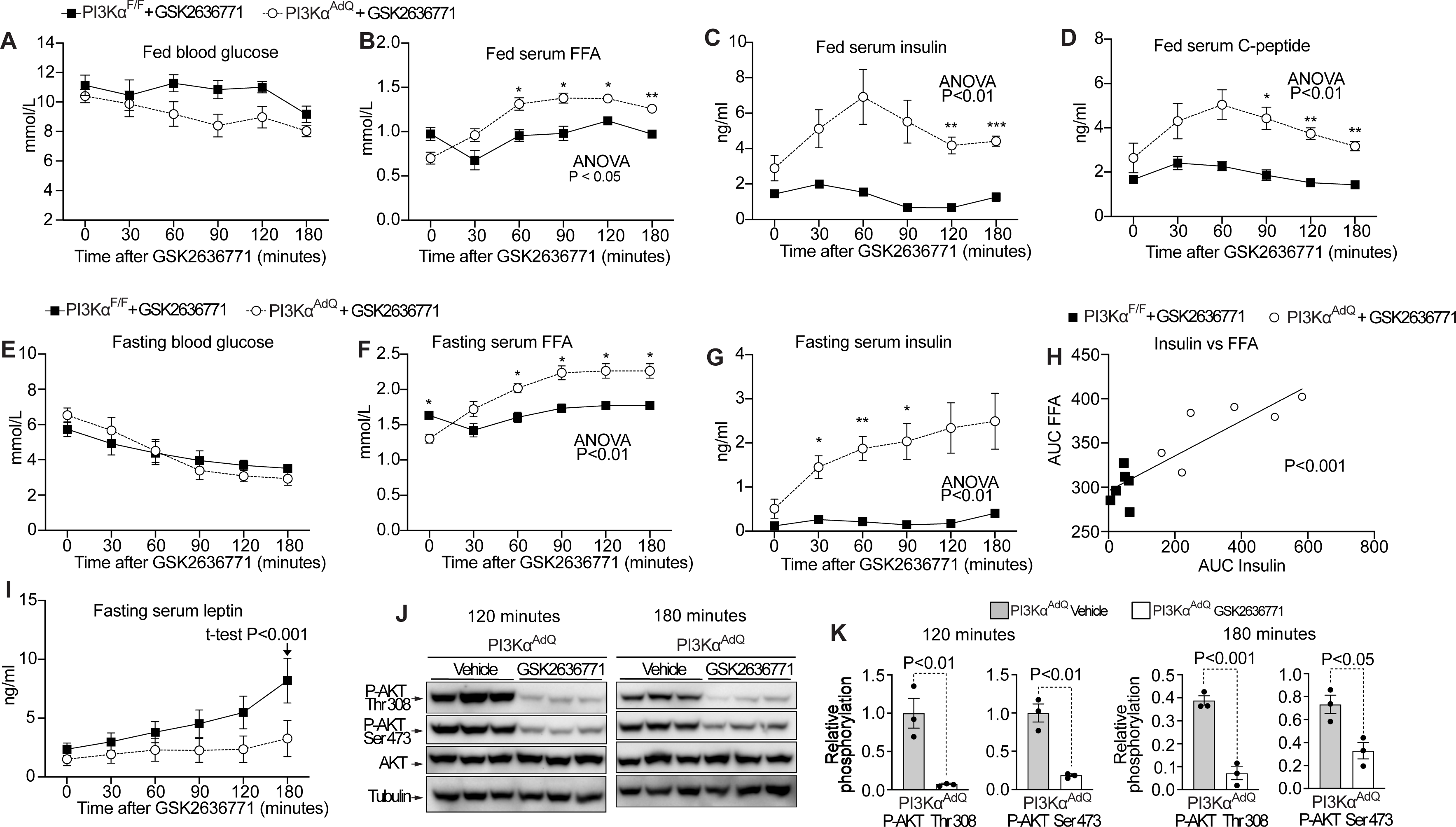
The adipoincretin effect operates in fed and fasted mice independently from glucose administration. (A) Blood glucose of fed PI3Kα^AdQ^ and PI3Kα^F/F^ mice injected with 6 mg/Kg of GSK2636771. (B) Serum levels of FFA; (C) serum insulin; and (D) serum C-peptide of the mice from A. (E) Blood glucose of mice as from A but injected with 6 mg/Kg of GSK2636771 after 12 hours of fasting. (F) Serum levels of FFA; and (G) serum insulin of the mice from E. (H) Simple linear regression of area under the curves (AUC) of insulin vs FFA; and (I) leptin of the mice from E. (J) Immunoblot analysis of AKT phosphorylation in epididymal adipose tissue from PI3Kα^AdQ^ mice fasted for 12 hours and sacrificed 120 and 180 minutes after injection of 6 mg/Kg of GSK2636771. n=5 mice for A-D and n=6 mice for E-I, n=3 for J. Data are mean ± s.e.m. Statistical analysis was performed using repeated measures, two-way ANOVA for A-J, Šídák’s multiple comparisons test for A-G, and unpaired t-test for H.

The results in fasting conditions are particularly remarkable as in PI3Kα^AdQ^ mice starved overnight, serum insulin kept increasing after GSK2636771 injection despite blood glucose abundance being low and decreasing during the experiment (Figures 3E-G). One mouse showed 0.43 ng/ml of insulin and 6.1 mmol/l of glucose before injection and 4.3 ng/ml of insulin and 2.5 mmol/l of glucose 180 minutes post-injection. The fact that the adipoincretin effect can operate in hypoglycemic mice with decreasing blood glucose, demonstrates that the adipoincretin effect is not driven by increased blood glucose. By contrast, we have found a strong linear correlation between the area under the curve (AUC) of serum FFA and serum insulin, showing that the adipoincretin effect is closely associated with lipolysis (Figure 3H).

Leptin is an insulin-regulated adipokine (Saladin et al., 1995), which controls adipostasis and insulin secretion (Coleman and Hummel, 1974; Covey et al., 2006). Therefore, we measured serum leptin levels in PI3Kα^F/F^ and PI3Kα^AdQ^ fasted mice after GSK2636771 injection. Compared to PI3Kα^F/F^ mice, PI3Kα^AdQ^ mice showed reduced leptin levels (Figure 3I). However, this difference in serum leptin was significant only 180 minutes after the GSK2636771 injection, indicating that the adipoincretin effect is not initiated by reduced serum leptin.

To further learn about the dynamic of PI3K inhibition, we have measured adipose tissue AKT phosphorylation at late time points, 120 and 180 minutes after GSK2636771 injection in fasted PI3Kα^AdQ^ mice. The results show a sustained inhibition of AKT phosphorylation (Figures 3J and K).

We conclude that the adipoincretin effect operates in fed and fasting conditions over a broad range of blood glucose concentrations, it is not initiated by reduced serum leptin and is associated with reduced adipocyte PI3K-AKT signaling and increased FFA.

### The adipoincretin effect is not associated with glucogenic or insulinogenic amino acids or exacerbated glucose counterregulatory response

We have measured the abundances of gluconeogenic and insulinogenic amino acids by ultra-performance liquid chromatography (UPLC) coupled with mass spectroscopy (MS), and of the hormones implicated in the glucose counterregulatory response by ELISA. PI3Kα^AdQ^ mice and littermate control PI3Kα^F/F^ mice were fasted overnight and sacrificed 90 minutes after the GSK2636771 injection. Plasma was collected at time point 0 (before injection) and 90 minutes post-injection. As expected, GSK2636771 markedly increased plasma C-peptide in PI3Kα^AdQ^ mice but not in control PI3Kα^F/F^ mice (Figure 4A). Blood glucose was slightly reduced during the experiment to a similar extent in the two genotypes (Figure 4B). PI3Kα^AdQ^ mice and littermate control PI3Kα^F/F^ mice also showed similar abundances of all the amino acids tested (Figure 4C). The plasma concentrations of adrenaline and corticosterone were increased during the experiment, which is consistent with a stress response to the experimental manipulation. However, plasma concentrations of adrenaline and corticosterone were virtually identical in PI3Kα^AdQ^ mice and PI3Kα^F/F^ control mice (Figures 4D and E). We have observed a trend for glucagon to increase in PI3Kα^AdQ^ mice at 90 minutes, but this trend was quantitatively marginal and statistically not significant (Figure 4F). Growth hormone was undetectable in plasma during the experiments (Figure 4G), which is consistent with a recent study showing that growth hormone levels are drastically reduced in mice fasted overnight (de Sousa et al., 2023).

**Figure 4.**
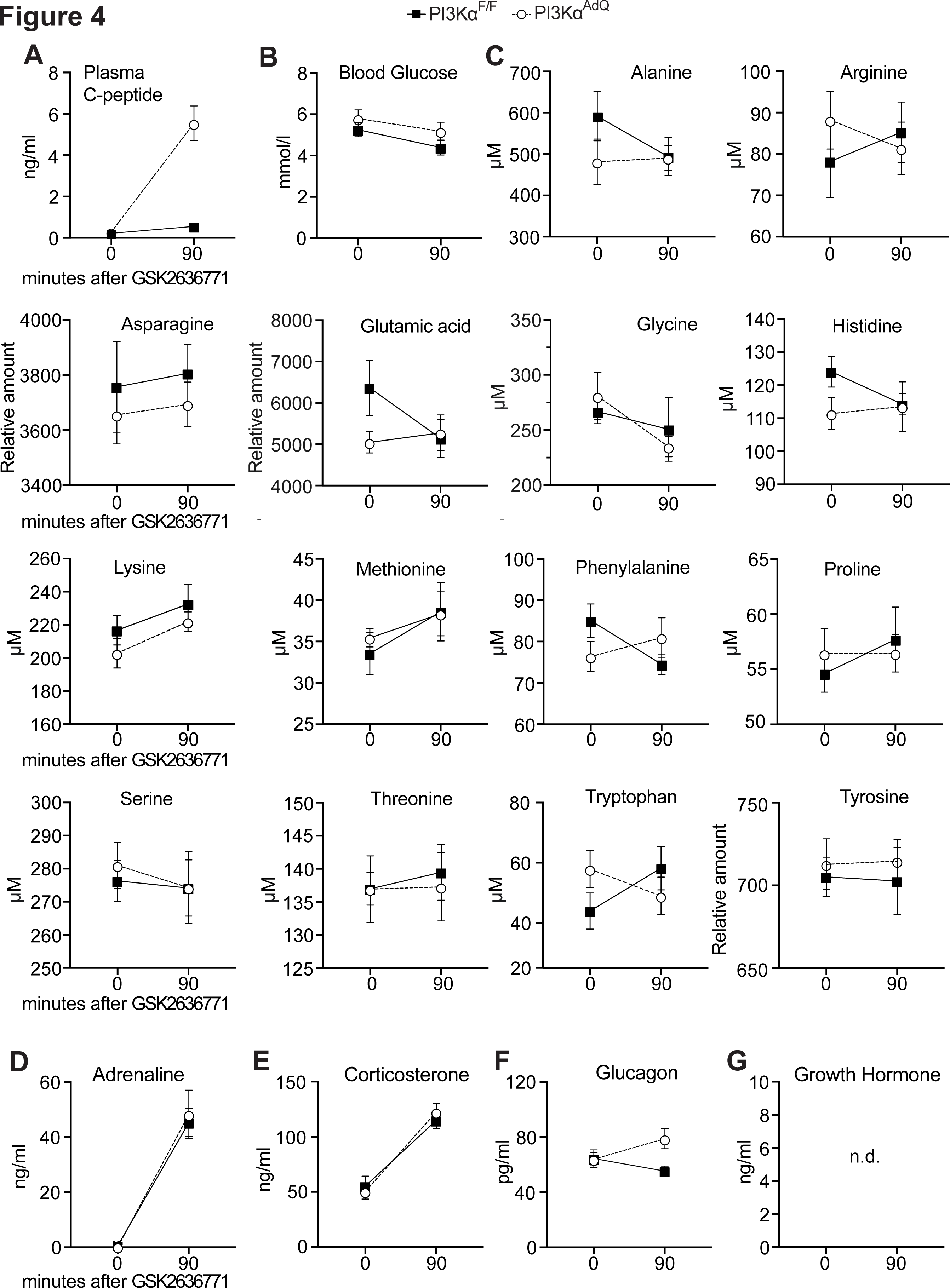
Amino acids and counterregulatory hormones in the adipoincretin effect (A) Plasma abundance of C-peptide in 12 hours fasted PI3Kα^AdQ^ and PI3Kα^F/F^ mice 90 minutes after injection of 6 mg/kg of GSK2636771. (B) Blood glucose of the mice in A. (C) plasma abundance of different gluconeogenic and insulinogenic amino acids. Plasma levels of (D) adrenaline, (E) corticosterone, (F) glucagon, and (G) growth hormone were measured by ELISA. n= 8 mice. Data are mean ± s.e.m. Statistical analysis was performed using repeated measures two-way ANOVA.

Overall, the adipoincretin effect was not associated with an increased abundance of gluconeogenic or insulinogenic amino acids or with hormones mediating the glucose counterregulatory response.

### The adipoincretin effect is associated with reduced plasma BCAA and increased 3OH- 4C-carnitine

We have investigated the role of candidate adipokines and metabolites in the adipoincretin effect. PI3Kα^F/F^ mice and PI3Kα^AdQ^ mice were fasted overnight and sacrificed 30 or 90 minutes after the GSK2636771 injection. Compared to baseline levels, plasma abundance of C-peptide was substantially increased 30 and 90 minutes after GSK2636771 injection in PI3Kα^AdQ^ mice but not in PI3Kα^F/F^ control mice (Figures 5A and S5A). Blood glucose levels were similar in PI3Kα^F/F^ and PI3Kα^AdQ^ mice 30 minutes after GSK2636771 injection and slightly, but significantly, reduced in PI3Kα^AdQ^ mice 90 minutes after GSK2636771 injection (Figures 5B and S5B). We measured plasma abundances of the adipokines Adipsin, FABP4, and Asprosin, which have been implicated in the control of fasting plasma insulin (Lo et al., 2014; Romere et al., 2016; Scheja et al., 1999). The plasma abundances of these adipokines were similar in PI3Kα^F/F^ and PI3Kα^AdQ^ mice at 30 and 90 minutes after GSK2636771 (Figure S6).

**Figure 5.**
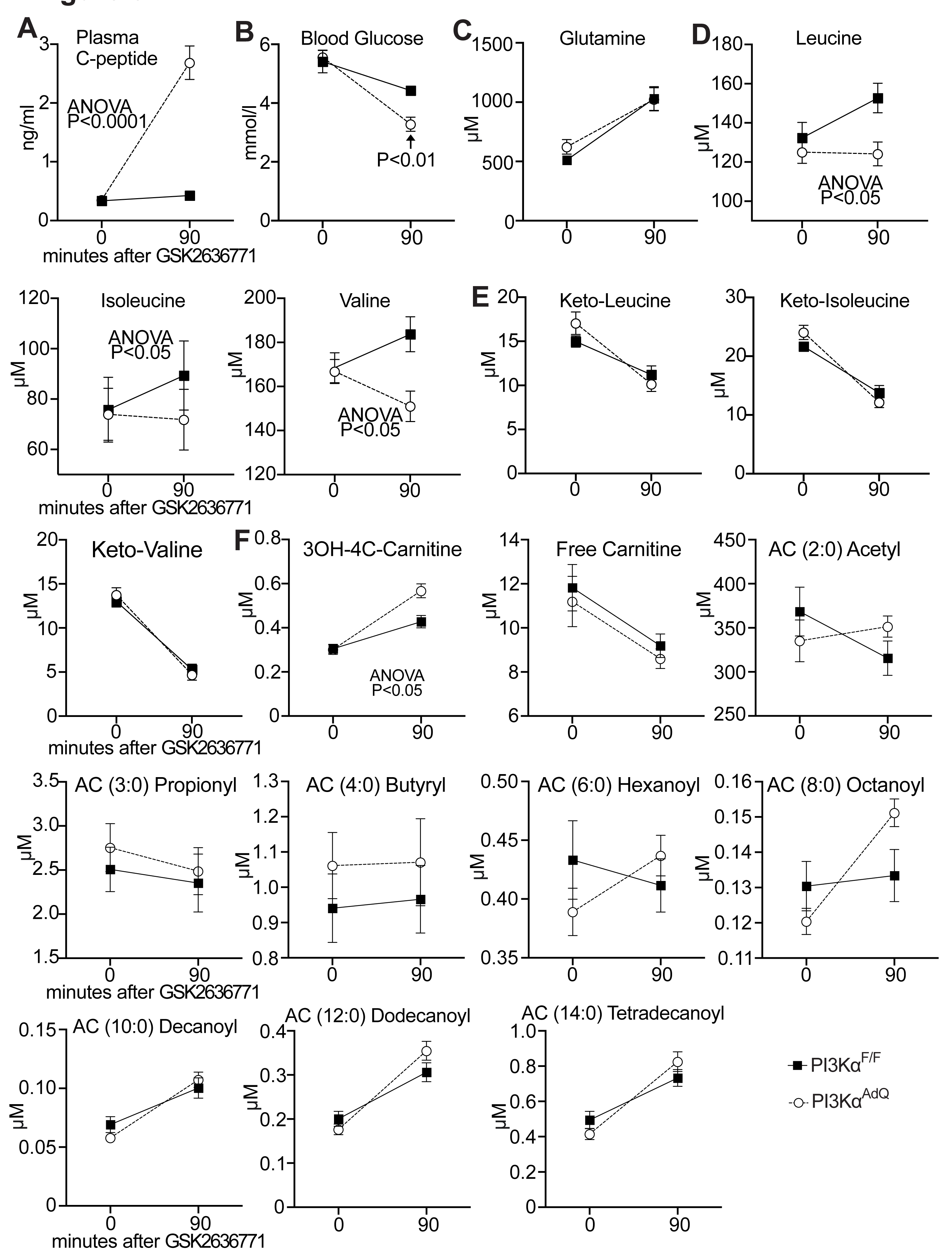
The adipoincretin effect is associated with reduced BCAA and increased 3OH-4C-Carnitine. (A) Plasma abundance of C-peptide in 12 hours fasted PI3Kα^AdQ^ and PI3Kα^F/F^ mice after injection of 6 mg/kg of GSK2636771. (B) Blood glucose of the mice in A. (C) Plasma abundance of glutamine; (D) branched-chain amino acids (BCAA) and (E) branched-chain ketoacids (BCKA) from the mice in A. (F) Plasma levels of free carnitine and acyl-carnitines in the mice from above. n= 10 mice. Data are mean ± s.e.m. Statistical analysis was performed using repeated measures two-way ANOVA and Šídák’s multiple comparisons test for B.

The branched-chain amino acid (BCAA) leucine, in combination with glutamine, potently induces insulin secretion in cultured β-cells (Sener and Malaisse, 1980). PI3Kα^F/F^ and PI3Kα^AdQ^ mice showed similar plasma glutamine levels at all time points. Still, plasma BCAA abundances were significantly reduced in PI3Kα^AdQ^ mice compared to PI3Kα^F/F^ control mice 90 minutes after GSK2636771 injection (Figures 5C, D and S5C, D). Plasma abundances of branched-chain ketoacids (BCKA) were virtually identical in PI3Kα^F/F^ and PI3Kα^AdQ^ mice (Figures 5E and S5E).

To further learn about the effects of adipocyte PI3K inhibition on fatty acids metabolism, we measured the abundance of acyl-carnitines in the plasma of the mice above. Compared to control PI3Kα^F/F^ mice, PI3Kα^AdQ^ mice showed an increased abundance of plasma 3-hydroxybutyrylcarnitine (3OH-4C-Carnitine) 90 minutes after GSK2636771 injection (Figure 5F). Other acylcarnitines and free carnitine were not significantly altered (Figures 5F and S5F).

These results indicate that the adipoincretin effect is not associated with increased plasma abundances of Adipsin, FABP4, Asprosin, glutamine, or BCKA but is associated with reduced BCAA and increased 3OH-4C-Carnitine.

### Contribution of lipolysis and FFA to the adipoincretin effect

β-adrenergic agonists induce lipolysis and stimulate insulin secretion in-vivo (Imura et al., 1971; Porte, 1967; Sennitt et al., 1985), whereas inhibition of lipolysis with nicotinic acid (NA) reduces insulin secretion in fasting (Balasse and Ooms, 1973; Boden et al., 1998; Dobbins et al., 1998a; Dobbins et al., 1998b; Stein et al., 1996).

To investigate the role of lipolysis in the adipoincretin effect, PI3Kα^AdQ^ mice were fasted overnight and injected with 6 mg/kg of GSK2636771 alone or GSK2636771 with 56 mg/kg of the adipose tissue triglyceride lipase (ATGL) inhibitor Atglistatin. As expected, GSK2636771 markedly increased insulin secretion over the baseline of PI3Kα^AdQ^ mice. However, Atglistatin cotreatment significantly reduced the effects of GSK2636771 on serum FFA and insulin secretion in PI3Kα^AdQ^ mice (Figures 6A-F). Notably, the Atglistatin treatment was associated with a trend toward increased blood glucose (Figures 6A and E). However, the AUC of serum insulin did not show a linear correlation with the AUC of blood glucose or of FFA (Figures 6G and H).

**Figure 6.**
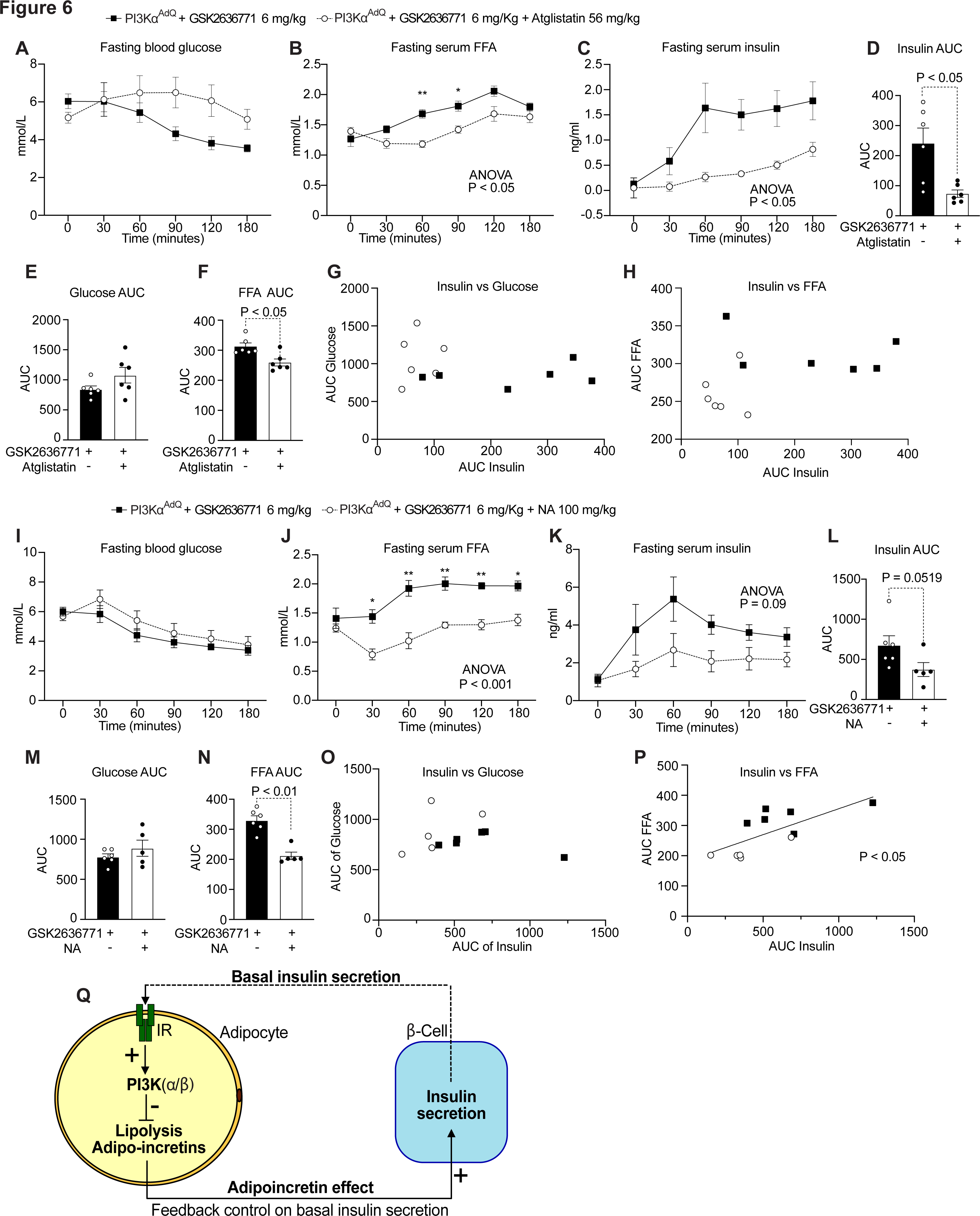
Contribution of lipolysis and FFA to the adipoincretin effect. (A) Blood glucose of 12 hours fasted PI3Kα^AdQ^ mice after injection of 6 mg/kg of GSK2636771 in the presence or absence of 56 mg/kg of Atglistatin. (B) The abundance of serum FFA and of (C) serum insulin from the mice in A. (D) area under the curve (AUC) of serum insulin in C. (E) AUC of fasting blood glucose levels in A; (F) AUC of FFA levels in B. (G) Simple linear regression of AUCs of insulin vs glucose; (H) Simple linear regression of AUCs of insulin vs FFA. (I) Blood glucose of 12 hours fasted PI3Kα^AdQ^ mice after injection of 6 mg/kg of GSK2636771 in the presence or absence of 100 mg/kg of nicotinic acid. (J) Serum FFA and (K) serum insulin from the mice above. (L) AUC of fasting insulin from K. (M) AUC of fasting blood glucose levels in I; (N) AUC of FFA levels in J. (O) Simple linear regression of the AUCs of insulin vs glucose. (P) Simple linear regression of AUCs of insulin vs FFA. (Q) During feeding, insulin secretion is governed mainly by the glucoincretin effect, which requires increased blood glucose. During fasting, the glucoincretin effect is absent, and blood glucose concentrations are minimal. Our findings indicate that baseline insulin secretion is mainly governed by closed-loop negative feedback mediated by insulin-PI3K signaling in the adipocyte. The system can be regarded as an adipostat, a “cruise control” for lipolysis where adipocyte PI3K activity links lipolysis with basal insulin secretion through the adipoincretin effect. n=6 mice for A-H, for I-P: n=6 for GSK2636771 and n=5 for GSK2636771 with NA. Data are mean ± s.e.m. Statistical analysis was performed using repeated measures two- way ANOVA, Šídák’s multiple comparisons test, Mann-Whitney for AUC, and simple linear regression for the relationship analysis of two AUC.

We next investigated the effects of nicotinic acid (NA), which inhibits lipolysis by reducing cAMP in adipocytes, on the adipoincretin effect. PI3Kα^AdQ^ mice were fasted overnight and injected with either 6 mg/kg of GSK2636771 alone or GSK2636771 with 100 mg/kg of NA. Blood glucose levels were similar between groups and decreased during the experiment (Figure 6I). Serum FFA abundance was increased over the baseline by GSK2636771, but this increase was completely blocked by the NA cotreatment (Figure 6J). GSK2636771 increased serum insulin levels in PI3Kα^AdQ^ mice cotreated with NA (Figure 6K). We observed a substantial trend, although not statistically significant, toward a reduced adipoincretin effect in PI3Kα^AdQ^ mice treated with GSK2636771 in the presence of NA (Figures 6K and L). Furthermore, AUC of serum insulin showed a significant linear correlation with the AUC of serum FAA, but not of blood glucose (Figures 6O and P).

Overall whereas we have observed a clear association between lipolysis and the adipoincretin effect, these results also indicate that the adipoincretin effect cannot be easily explained by a general increase of circulating FFA alone. Indeed, Atglistatin was less effective than NA in reducing serum FFA but more effective than NA in reducing serum insulin (Figures 6B,C and J,K), without a correlation between the AUCs of serum FFA and insulin (Figure 6H).

Although for the GSK2636771-NA cotreatment experiment there was a linear correlation between the AUCs of serum FFA and insulin (Figure 6P), we also observed at 30 minutes post-injection the opposite change in serum FFA abundance (reduced) and insulin secretion (increased) in PI3Kα^AdQ^ mice cotreated with GSK2636771 and NA (Figures 6J and K). This finding was reproduced in an independent cohort of PI3Kα^AdQ^ mice cotreated with GSK2636771 and NA, which showed an even more pronounced dissociation between serum FFA and serum insulin (Figures S7A-D).

In conclusion, whereas these results are consistent with a functional role for adipocyte lipolysis in the adipoincretin effect, also show that a general increase in circulating FFA alone cannot fully explain this phenomenon and indicates that PI3K activity in the adipocyte regulates another signal contributing to the adipoincretin effect (e.g. an adipoincretin).

### The hyperinsulinemia induced by systemic PI3K inhibition can be reduced with NA

The hyperinsulinemia associated with systemic PI3K inhibition reduces the therapeutic index of PI3K inhibitors in cancer therapy (Hopkins et al., 2018; Vasan and Cantley, 2022).

Therefore, we have investigated the contribution of lipolysis and increased FFA to the insulin secretion induced by whole-body PI3K inhibition (which also increases blood glucose).

Wild-type (WT) C57Bl6/J mice were fasted overnight and injected intraperitoneally with Taselisib (3.5 mg/kg) with or without Atglistatin (56 mg/kg). As expected blood glucose and serum FFA abundances were increased in mice treated with Taselisib. However, while the cotreatment with Atglistatin partially reduced serum FFA, also markedly exacerbated hyperglycemia (Figures 7A, and B). Nevertheless, compared to mice treated with Taselisib only, mice cotreated Atglistatin showed reduced serum insulin during the first 60 minutes after injection and similar serum insulin at later time points despite substantially higher blood glucose (Figure 7C). There was no linear correlation between AUC of insulin and AUC of either glucose or FFA (Figures 7D-H).

**Figure 7.**
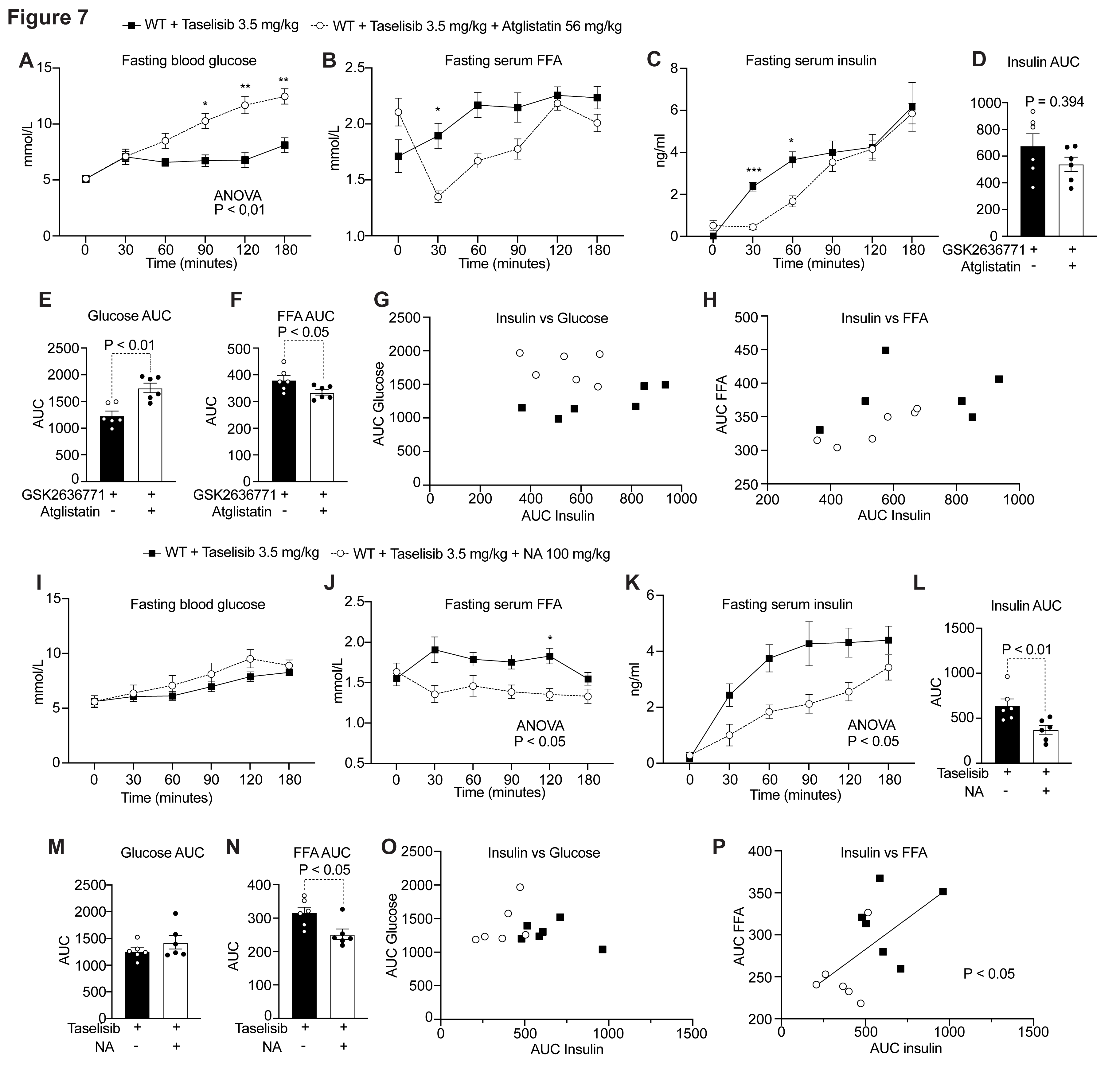
Contribution of lipolysis to systemic PI3K inhibition (A) Blood glucose of 12 hours fasted wild-type C57Bl/6 mice after injection of 3.5 mg/kg of Taselisib in the presence or absence of 56 mg/kg of Atglistatin. (B) Serum FFA and (C) serum insulin from the mice above. (D) Area under the curve (AUC) of serum insulin in C. (E) AUC of fasting blood glucose levels in A; (F) AUC of serum FFA in B. (G) Simple linear regression of the AUCs of insulin vs glucose. (H) Simple linear regression of the AUCs of insulin vs FFA. (I) Blood glucose of 12 hours fasted C57Bl/6 wild-type mice after injection of 3.5 mg/kg of Taselisib in the presence or absence of 100 mg/kg of nicotinic acid (NA). (J) Serum FFA and (K) serum insulin from the mice above. (L) AUC of serum insulin from K. (M) AUC of fasting blood glucose in I; (N) AUC of serum FFA levels in J. (O) Simple linear regression of the AUCs of insulin vs glucose. (P) Simple linear regression of the AUCs of insulin vs FFA. n=6 mice. Data are mean ± s.e.m. Statistical analysis was performed using repeated measures two-way ANOVA, Šídák’s multiple comparisons test, Mann-Whitney for AUC, and simple linear regression for the relationship analysis of two AUC.

We have investigated the effects of NA on systemic PI3K inhibition by comparing WT C57Bl6/J mice fasted overnight and injected with either 3.5 mg/kg of Taselisib alone with mice injected with Taselisib and NA (100 mg/kg).

WT mice injected with of Taselisib alone showed increased blood glucose, serum FFA, and substantially elevated serum insulin (Figures 7I-K). Cotreatment with NA completely prevented the increase of serum FFA without affecting blood glucose (Figures 7I and J).

Taselisib increased serum insulin also in the presence of NA (Figure 7K). However, NA significantly reduced the increase of serum insulin induced by Taselisib (Figure 7K). AUC of serum insulin showed a linear correlation with AUC of FFA but not AUC of blood glucose.

These findings were reproduced in an independent cohort of mice in fasting and fed conditions, with the only difference that there was a linear correlation also between the AUC of insulin and blood glucose (Figures S7E-L and R, S). We tested the effects of NA also with another pan-PI3K inhibitor, Buparlisib. However, at the dose tested (8 mg/kg), Buparlisib caused only a modest increase in serum insulin and blood glucose without increasing serum FFA (Figures S7M-O). Buparlisib NA cotreatment showed a trend toward reduced insulin (Figures S7M-O). AUC of serum insulin was associated with AUC of blood glucose but not FFA (Figures S7P, T).

In conclusion our results show that the hyperinsulinemia caused by systemic PI3K inhibition can be significantly reduced by NA coadministration.

## DISCUSSION

We now show that, during fasting, adipocyte lipolysis and basal insulin secretion are coregulated by redundant PI3Kα and PI3Kβ activities within the adipocyte independently from blood glucose through a feedback signal that we named the adipoincretin effect.

Our understanding of insulin-PI3K signaling in the adipocyte in-vivo has been limited by the fact that genetic ablation of the InsR or AKT1 and AKT2 in mice adipocytes leads to severe lipoatrophy with no functional adipose tissue left to study its physiological function (Boucher et al., 2016; Qiang et al., 2016; Sakaguchi et al., 2017; Shearin et al., 2016). To overcome this issue, we first discovered that adipocyte insulin signaling in mice is mediated by the redundant activity of PI3Kα and PI3Kβ, and then we exploited this redundancy to achieve an adipose tissue-selective pharmacological inhibition of PI3K activity. Acute adipocyte-selective PI3K inhibition, achieved by administering GSK2636771 to PI3Kα^AdQ^ mice, blocked the effect of glucose on lipolysis and increased serum FFA in mice fasted overnight. This finding substantially advances our understanding of insulin action on lipolysis, showing that adipocyte PI3K is active even after prolonged fasting to control the rates of lipid mobilization.

Most importantly, we discovered that acute inhibition of PI3K activity in adipocytes caused a substantial and sustained increase in insulin secretion. This phenomenon, the adipoincretin effect, is independent of the feeding state. However, because the adipoincretin effect is triggered by low adipocyte PI3K activity and is associated with elevated lipolysis, it is logical to deduce that the physiological function of the adipoincretin effect is to control the rate of lipid mobilization during fasting. Food-induced insulin secretion is mainly governed by the action of glucoincretins in synergy with increased blood glucose and, to a lesser extent, by the cephalic phase (Berthoud et al., 1981; Nauck et al., 1986). By contrast, our results show that basal insulin secretion during fasting is chiefly controlled by the adipoincretin effect, which is inhibited by adipocyte PI3K signaling (Figure 6Q).

This concept stands in stark contrast to the current paradigm of the insulin system, which focuses on glycemic control and indicates that, during fasting, the insulin system operates as a “cruise control” for lipolysis, a true adipostat (Figure 6Q) (Solinas and Becattini, 2023).

It has been reported that adipocyte insulin signaling controls hepatic glucose production after prolonged fasting when liver glycogen is depleted (Perry et al., 2015; Petersen and Shulman, 2018; Titchenell et al., 2017; Titchenell et al., 2016). However, our data show that the adipoincretin effect can be dissociated from increased blood glucose. The effect of adipocyte PI3K inhibition on insulin secretion was far more potent than the effect on insulin secretion caused by an intraperitoneal injection of 2 g/kg of glucose (Figures 2A-B). In fasted mice, the adipoincretin effect was induced while blood glucose decreased from more than six mmo/L to about three mmol/L (Figures 3E and G). Therefore, the adipoincretin effect cannot be explained by an increase even transient in blood glucose.

Activation of lipolysis by β-adrenergic agonists is known to induce insulin secretion while reducing blood glucose (Grujic et al., 1997; Imura et al., 1971; Porte, 1967; Yoshida, 1992). The adipoincretin effect was clearly associated with increased lipolysis (Figure 3 E-H and Figure 6). However, we could partially dissociate the adipoincretin effect from increased FFA by co-treating PI3Kα^AdQ^ mice with the PI3Kβ-selective inhibitor GSK2636771 in the presence of NA or by using the less potent PI3Kβ-selective inhibitor TGX221 (Figures S3A-B and S7A-D). This observation is consistent with studies showing that increasing circulating FFA by intralipids and heparin infusion does not substantially increase insulin secretion (Amery et al., 2000; Balent et al., 2002; Becattini et al., 2023; Grujic et al., 1997; Hargreaves et al., 1991; Schenk et al., 2005; Smiles et al., 2019; Yoshida, 1992). Atglistatin showed a more pronounced inhibition of the adipoincretin effect than NA despite a partial reduction of FFA, and without correlation between FFA AUC and insulin AUC (Figure 6). Finally 30 minutes after NA injection serum insulin was increased while serum FFA were reduced (Figures 6J, K and S7B, C).

Collectively, these pieces of evidence indicate that a general increase of serum FFA alone is not sufficient to fully explain the adipoincretin effect, which likely depends on a specific adipocyte-derived “adipoincretin” hormone whose release is controlled by PI3K activity.

Our data exclude a role for altered amino acid metabolism. Specific adipokines could mediate the glucoincretin effect, and leptin is a logical candidate (Ahima et al., 1996; Coleman and Hummel, 1974; Covey et al., 2006; D’Souza A et al., 2016; Saladin et al., 1995; Trayhurn et al., 1995). Compared to control mice, mice with adipocyte-selective PI3K inhibition showed reduced serum leptin, but this difference was significant only by the end of the experiment, after the induction of insulin secretion (Figure 3). We conclude that the adipoincretin effect is not initiated by the reduced abundance of circulating leptin. However, low circulating leptin abundance likely drives insulin secretion after a prolonged period of starvation when the adipose tissue is progressively depleted. This view is supported by a study showing that inducible ablation of the insulin receptor in adipocytes of adult mice within a few days causes severe lipoatrophy and hypoleptinemia in association with hyperinsulinemia, which can be prevented by continuous leptin infusion (Sakaguchi et al., 2017).

We investigated other candidates “adipoincretins”: Adipsin, FABP4, and Asprosin (Lo et al., 2014; Romere et al., 2016; Scheja et al., 1999). Adipocyte-specific PI3K inhibition did not affect the circulating levels of Adipsin, FABP4, and Asprosin. However, it remains to be found whether adipocyte PI3K signaling plays a role in controlling the activity of one or more of these adipokines.

We have found that the plasma abundance of 3OH-4C-Carnitine was significantly elevated in mice 90 minutes after adipocyte-selective PI3K inhibition. 3OH-4C-Carnitine is elevated in humans affected by a form of hyperinsulinism caused by loss of function mutations on the gene encoding for L-3-hydroxyacyl-CoA-dehydrogenase (HADH) (Clayton et al., 2001).

However, the role of adipocyte HADH activity in insulin secretion remains unresolved, and more research is needed to understand the significance of increased 3OH-4C-Carnitine on insulin secretion. Some insight may be derived from our data on ATGL inhibition. The fact that Atglistatin was more effective in reducing the adipoincretin effect than NA, without showing a linear correlation between AUCs of serum insulin and FFA suggests that the adipoincretin effect may depend on a specific enzymatic activity of ATGL. Indeed, besides its lipase activity, ATGL was implicated in the synthesis of branched fatty acid esters of hydroxy fatty acids (FAHFAs) (Patel et al., 2022). Further research in this direction is granted.

The hyperinsulinemia induced by PI3K inhibitors dampens the efficacy of PI3K- targeted cancer therapy (Vasan and Cantley, 2022). We now show that inhibition of PI3K in adipocytes induces insulin secretion independently from blood glucose and that NA significantly reduces the hyperinsulinemia induced by the pan PI3K inhibitor Taselisib.

Future research will test whether NA (vitamin B3) supplementation can improve the efficacy of PI3K inhibitors in cancer therapies.

It has been proposed that in obese subjects with normal glucose tolerance, hyperinsulinemia can be explained mainly by increased baseline insulin secretion at fasting (Polonsky et al., 1988). Because adipose tissue lipolysis and fasting serum insulin are associated in obese subjects (Fryk et al., 2021), the development of hyperinsulinemia and insulin resistance in obesity may be driven by an exacerbated adipoincretin effect due to increased fat mass.

Future research on the adipoincretin effect holds great promise for developing targeted therapies for cancer, obesity, and type-2 diabetes.

## Limitations of the study

The lack of identification of the specific adipoincretin endocrine signals is the major limitation of our study. However, this limitation does not make the discovery of the adipoincretin effect any less solid or disruptive. By analogy, the finding that GLP-1 is a glucoincretin was published 23 years after demonstrating the glucoincretin effect, a disruptive discovery that transformed our understanding of how food induces insulin secretion (Rehfeld, 2018). The discovery of the adipoincretin effect transforms our understanding of how the insulin system operates, revealing that during fasting, the insulin system operates as an adipostat (Figure 6Q) (Solinas and Becattini, 2023). This model has a major impact on obesity, diabetes, and cancer research.

## ACKNOWLEDGMENTS

We are grateful to Prof. Claes Wollheim at Lund University for our conversations on β-cell biology and to Prof. Paolo De Los Rios at the EPFL for our conversations on the properties of feedback control systems. This study is supported by grants from the Novo Nordisk Foundation NNF19OC0057174; the Swedish Research Council 2022-01033; the Cancerfonden 20 0840 PjF; and the Diabetesfonden DIA2022-753 to G.S.

## AUTHOR CONTRIBUTIONS

B.B codesigned and performed most experiments, interpreted data, and contributed to writing the manuscript; A.M generated the PI3Kα^AdQ^ mice and characterized the PI3Kα deletion; M.H. and J.B. performed targeted metabolomics; G.S. conceived the study, analyzed, and interpreted the results, and wrote the first version of the manuscript. All authors have read and approved the manuscript.

## STAR*METHODS

### KEY RESOURCES TABLE

| REAGENT or RESOURCE | SOURCE | IDENTIFIER |
| --- | --- | --- |
| **Antibodies** |  |  |
| AKT Phospho (Thr308) | Cell Signaling | 4056 |
| AKT Phospho (Ser373) XP | Cell Signaling | 4060 |
| AKT | Cell Signaling | 9272 |
| Erk 1/2 (p44/42 MAPK)<br>Phospho(Thr202/Tyr204) XP | Cell Signaling | 9101 |
| Erk 1/2 (p44/42 MAPK) | Cell Signaling | 9102 |
| IRS1 | Millipore | 06248 |
| IRS2 | Millipore | MAB515 |
| PI3K p85 | Cell Signaling | 4257S |
| PI3K p85α | Abcam | Ab133595 |
| PI3K p55γ | Abcam | Ab238509 |
| Tubulin $\alpha/\beta$ | Cell Signaling | 2148S |
| PI3Kα | Cell Signaling | 4249 |
| PI3Kβ | Cell Signaling | 3011 |
| PI3Kδ | Millipore | 04-401 |
| PI3Kγ | a gift of Prof. Matthias Wymann<br>University of Basel | (Russian) |
| HSL Phospho (Ser563) | Cell Signaling | 4135 |
| HSL Phospho (Ser660) | Cell Signaling | 4126 |
| HSL Phospho (Ser565) | Cell Signaling | 4137 |
| HSL | Cell Signaling | 4107 |
| **Chemicals, Peptides, Recombinant Proteins and Inhibitors** |  |  |
| Insulin | Humalog Lilly | VL7510 |
| Glucose | Acros | 410955000 |
| BSA | Fisher Scientific | BP9702 |
| ECL-anti rabbit IgG HRP | GE Healthcare | NA934V |
| ECL-anti mouse IgG HRP | GE Healthcare | LNA931V |
| Collagenase P | Roche | 11213865001 |
| GSK2636771 (Inhibitor PI3Kβ) | SelleckChem | S8002 |
| TGX-221 (Inhibitor PI3Kβ) | MedChem Express | HY-10114 |
| IPI549 (Inhibitor PI3Kγ) | MedChem Express(SelleckChem) | HY-100716 |
| PIK294 (Inhibitor PI3Kδ) | MedChem Express | HY-10303 |
| Buparlisib (BKM120, panPI3K inhibitor) | MedChem Express | HY-70063 |
| Taselisib (GDC-0032, panPI3K inhibitor) | MedChem Express | HY-13898 |
| Nicotinic Acid | Sigma | 72309 |
| Atglistatin | MedChem Express | HY-15859 |
| Cremophor EL | Millipore | 238470 |
| **Critical Commercial Assays** |  |  |
| Insulin ELISA Kit | Crystal Chem | 90080 |
| C-peptide Elisa Kit | Crystal Chem | 90050 |
| FFA (NEFA-HR set) | Fuji Film | 436-91995, 434-91795<br>91096 (standard) |
| Leptin Elisa Kit | R&D | MOB00B |
| Adipsin Elisa Kit | R&D | DY5430-05 |
| FABP4 Elisa Kit | BioVendor | RD191036200R |
| Asprosin Elisa Kit | Abbexa | abx585287 |
| Adrenaline Elisa Kit | LDN | BA E-5100R |
| Corticosterone Elisa Kit | IDS | AC-14F1 |
| Glucagon Elisa Kit | Crystal Chem | 81518 |
| Mouse Growth factor Elisa Kit | Crystal Chem | 80587 |
| Immobilon Western Chemiluminescence HRP substrate | Millipore | WBKLS0500 |
| Hematoxylin Solution (Mayer's, Modified) | Abcam | AB220365 |
| Eosin | Sigma | HT110116 |
| **Experimental Models: Organisms/Strains** |  |  |
| WT | C57BL/6 background | Janvier Labs |
| PI3K$\alpha^{Flox}$ | Mixed background | Provided by Victoria Rotter Sopasakis |
| PI3K$\alpha^{AdQ}$ | Mixed background | Provided by Victoria Rotter Sopasakis |
| **Software and Algorithms** |  |  |
| Image Lab software (version 5.2.1) | Bio-Rad | http://www.bio-rad.com/en-us/product/image-lab-software |
| Graph Pad Prism 7.0 | Graph Pad Software | N/A |
| AxioVision Software | Imaging Software | Carl Zeiss Microscopy |
| **Deposited Data** |  |  |
| DOI: 10.17632/tgsgkjz28w.1 | Immunoblots images | N/A |

### CONTACT FOR REAGENT AND RESOURCE SHARING

Further information and requests for reagents may be directed to the Lead Contact, Giovanni Solinas

#### Mice and In Vivo Studies

All mouse studies were approved by the Ethics Committee on Animal Care Use in Gothenburg, Sweden. Only male mice were used in this study. Mice were maintained at the EBM specific pathogen free facility of the University of Gothenburg under a 12-hour light /12-hour dark cycles and at a room temperature of 22°C.

PI3Kα LoxP floxed mice (PI3Kα^F/F^) in 129/SvJ and C57BL/6J mixed background were previously described (Molinaro et al., 2019). PI3Kα^AdQ^ mice and littermate control PI3Kα^F/F^ mice were obtained by crossing Adipoq-Cre mice in pure C57BL/6J background (Jackson labs) with the PI3Kα^F/F^ mice from above. Wild-type (WT) C57BL/6J mice were obtained from Janvier Labs.

For the insulin-tolerance test (ITT), mice were fasted for 4 hours and injected intraperitoneally with 1 I.U. of insulin per Kg of body weight. Blood was collected from the tail at the indicated post-injection times, and blood glucose concentrations were measured using a glucometer (Contour Next, Ascensia). For the glucose tolerance test (GTT), mice were fasted for either 4 hours (GTT in Figure S2) or overnight for glucose-mediate suppression of lipolysis (Figure 2) and injected intraperitoneally with either 1g or 2 g of glucose per Kg of body weight respectively. Blood was collected from the tail at the indicated time points and used to measure blood glucose and prepare serum to measure hormones and metabolites. For in vivo injection of PI3K inhibitors, the indicated dose of the compound was injected intraperitoneally into eight to twelve-weeks-old PI3Kα^AdQ^ mice and PI3Kα^F/F^ littermate controls or C57BL/6 mice. All mice were treated acutely. For each experiment, mice were injected only once immediately after the 0- minute time point, and the experiment lasted for a maximum of 180 minutes. We used a low dose of isoform-selective PI3K inhibitors (e.g., 6 mg/Kg of body weight for GSK2636771) to ensure isoform selectivity during the relatively short time of the experiment (no longer than 180 minutes). This dose was determined in a previous study for TGX221, which we identified as the lowest dose that increased blood glucose in mice lacking PI3Kα in their hepatocytes but not in control mice (Molinaro et al., 2019). The dose of all other isoform-selective inhibitors was estimated relative to the dose of TGX221 we used and the IC50 of the specific PI3K inhibitor.

The injectable solutions of the PI3K inhibitors were freshly prepared as follows: PI3K inhibitor was first dissolved in pure DMSO, and to this master liquid were added one by one and in this order PEG300, Tween 80, and water. The solution was mixed after each addition. The final concentration for the single components was as follows: 5% DMSO, 30% PEG, and 1% Tween-80. Control ‘‘vehicle solution’’ was 5% DMSO, 30% PEG, 1% Tween-80, and water. For solutions including Atglistatin, the latter was added to a water fraction containing 0.5% of Cremophor EL and dissolved by means of ultrasonication. The water solution thus obtained was added to the DMSO/PEG300/Tween80 mixture. For solutions including nicotinic acid (NA), the NA powder was directly dissolved in the final mix at the desired concentration; the acidity was adjusted at a value of pH 7 by adding NaOH 1M to the solution. Blood glucose was measured at the indicated time points after injection and blood was collected for further measurements of serum insulin, serum FFA, serum leptin, and plasma levels of C-peptide, BCAA, BCKA, aminoacids, free and acyl-carnitines.

At sacrifice, mice were anesthetized with Isofluorane (Baxter KDG 9623) and euthanized by CO2 and bleeding.

#### Insulin, C-peptide, Leptin, counterregulatory factors, and adipokine measurements

Insulin and C-peptide levels in serum and plasma were measured using an ELISA kit from Crystal Chem. Serum leptin was measured by ELISA kit from R&D. Counterregulatory factor adrenaline corticosterone, glucagon, and growth hormone were measured using Elisa kits from LDN, IDS, and Crystal Chem respectively. Plasma Adipsin (Mouse Complement Factor D) was measured by quantitative sandwich enzyme immunoassay using kits obtained from R&D Systems, and adipocyte FABP4 abundance in plasma was measured using an ELISA kit from BioVendor. All ELISA assays were performed according to the instruction of the manufacturer.

#### Hematoxylin and eosin (H&E) staining

Samples of adipose tissue pads and liver were obtained from 13-week-old mice. The samples were fixed in 10% formalin, embedded in paraffin, cut into 5 μm sections, and stained with hematoxylin (Abcam) and eosin (Sigma). H&E stained sections were imaged at 20X magnification with an AxioImager M1 microscope (Zeiss, Germany).

#### Adipocyte size distribution

Images of two fields for six mice were analyzed in Fiji using the open source plugin adiposoft. Cell surface area was calculated in microns and classified based on area μm^2^.

#### Immunoblot Analysis

For analysis of protein phosphorylation, we noticed that stripping procedures frequently reduce the overall immunoblot quality and sporadically create artifacts. Thereby protein samples were loaded in parallel. For controls on total proteins, tubulin was analyzed from the same membrane when possible. resolved by SDS-PAGE electrophoresis and then transferred to a PVDF membrane. The membrane-blocking procedure was carried out at room temperature in phosphate saline buffer (PBS) with 3% bovine serum albumin (BSA), and 0.3% tween solution (Cell Metabolism 30, 1400–1409.e1–e5, June 4, 2019 e4). The primary antibody was incubated in a cold room overnight in PBS with 0.3% tween and 3% BSA; the membranes were then washed three times with a PBS 0.3% tween solution and incubated with a horseradish peroxidase-conjugated secondary antibody (GE Healthcare) in PBS 0.3% tween 3% BSA for one hour at room temperature. After three washes in a PBS 0.3% tween solution and incubation with a detection reagent (Millipore), the signal was acquired using the Biorad ChemiDoc apparatus.

#### Analysis of plasma branched-chain amino acids and branched-chain keto acids

25 µl plasma and standard was precipitated with 250 µl methanol containing 1 µM of deuterated internal standards (Valine-^13^C5,^15^N, Leucine d3, Isoleucine d10, keto-leucine d7 and keto- isoleucine). After 10 minutes of vortex at 1400 rpm, the samples were centrifuged at 4000g for 10 minutes. 25 µl was removed and evaporated under a stream of nitrogen gas. The samples were reconstituted in 125 µl water and injected into the UPLC-MS/MS system. Separation was performed on a Waters Acquity BEH C8 column (2.1 x 100mm 1.7 µm) with a gradient consisting of water and acetonitrile, both with 0.1% formic acid. The detection was made on a Waters Xevo TQ-XS triple quadrupole mass spectrometer in MRM mode using intra-run polarity switching where the detection of amino acids was made in positive mode between 0-3 minutes. The polarity was then switched into a negative mode for the detection of the keto acids between 3-10 minutes. The standard curve containing all six amino- and keto acids was prepared in 20% methanol and treated the same way as the samples.

#### Analysis of plasma acylcarnitines and aminoacids

25 µl of plasma was precipitated with 250 µl of methanol containing 1 µM of stable isotope labelled internal standards (^1^ Alanine-D4, Ariginine-^13^C6 ^15^N5, Glutamic acid-D5, Glutamine-^13^C5 ^15^N2, Glycine-D5, Lysine-D4, Methionine-^13^C1 D3, Phenylalanine-D8, Proline-^13^C5 ^15^N1, Serine-^13^C3 D1 ^15^N1, Tryptophan-D5). After 10 minutes of vortex at 1400 rpm, the samples were centrifuged at 4000g for 10 minutes. 25 µl was removed and evaporated under a stream of nitrogen gas.

The samples were reconstituted in 125 µl of an acylcarnitine internal standard mix (NSK-B reference mix, Cambridge Isotope Laboratories) prepared in acetonitrile: methanol [3:1]. The samples were separated in HILIC mode on a Waters BEH Amide column (2.1 x 100 1.7µm) using 5 mM ammonium formate and 0.06% formic acid in 95% acetonitrile as mobile phase A and 10 mM ammonium formate and 0.12% formic acid in 100% Milli-Q water as mobile phase B. The detection was made on a Waters Xevo TQ triple quadrupole mass spectrometer in positive MRM mode. Quantification of acylcarnitines was made using a one-point calibration against the deuterated analog. For the acylcarnitines without a deuterated analog, the quantification was made against the deuterated acylcarnitine closest in retention time. Amino acids were quantified against an external calibration curve made with reference substances in 20% methanol. For some amino acids the reference substance was missing (asparagine, glutamic acid and tyrosine). For those amino acids no quantification was made. Instead the relative amount was attained by dividing the endogenous signal with the internal standard. For amino acids that lack the labelled analogue, another internal standard (with similar retention time) was used.

#### Glucose Stimulated insulin secretion (GSIS)

Primary mouse islet isolation was performed as previously described (Hugill A, Shimomura K, Cox RD. Islet Insulin Secretion Measurements in the Mouse. Curr Protoc Mouse Biol. 2016 Sep 1;6(3):256-271. doi: 10.1002/cpmo.14. PMID: 27584553). After isolation and overnight incubation at 37°C with humidified 5% CO2 air in RPMI supplemented with 1% P/S and 10% FBS, islets were transferred into a 24-well plate. 5 islets were hand-picked and transferred into each well in the presence of 200 μl low glucose buffer (Krebs Ringer Buffer KRB supplemented with 0.2% BSA and 2.8 mM glucose) and pre-incubated for 1 hour at 37°C. Islets underwent successive incubations for 20 minutes at 37°C with 200 μL of fresh KRB 0.2% BSA containing low glucose (2.8 mM), followed by high glucose (16.7 mM). At the end of each incubation, islets were transferred into a new well, and the supernatant was collected for analysis of insulin levels. At the end of the stimulation protocol, cells were lysed in 70 μL acid ethanol (75% absolute ethanol, 23.5% water, 1.5% HCl) and stored at 4°C overnight. Samples were dried with a SpeedVac and resuspended in 30 μL ultrapure water. DNA was quantified by Nanodrop (ThermoFisher, USA), and insulin was quantified using an ELISA kit (Crystal Chem) and normalized to DNA content.

### QUANTIFICATION AND STATISTICAL ANALYSIS

Data are expressed as means, and error bars indicate standard errors.

Only biological replicates, defined as data from tissues of different mice, are used for statistical analysis. The exact number of mice for a specific experiment is indicated in the figure legends. For all experiments, littermate male mice with similar body weight were used.

Sample size was determined empirically based on previous experience and practical considerations. In the time course after injection, a minimum of 5 mice was used; for all other in vivo studies, a minimum of 3 biological replicates was analyzed when parametric statistical tests were used, whereas a minimum of 4 biological replicates was analyzed when non-parametric statistical tests were used. In analyses where two different categorical variables are considered (GTT, ITT, mice weight gain, and time-course analyses), repeated measures (RM) two-way ANOVA was used. P values in the figures always refers to the group effect. The exact P values are shown when close to 0.05.

All statistical analysis was performed with GraphPad Prism software.

