## Supplemental figures for "Adipocyte PI3K Links Adipostasis with Basal Insulin Secretion Through an Adipoincretin Effect"

Seven supplemental figures.

**Fig. S1**

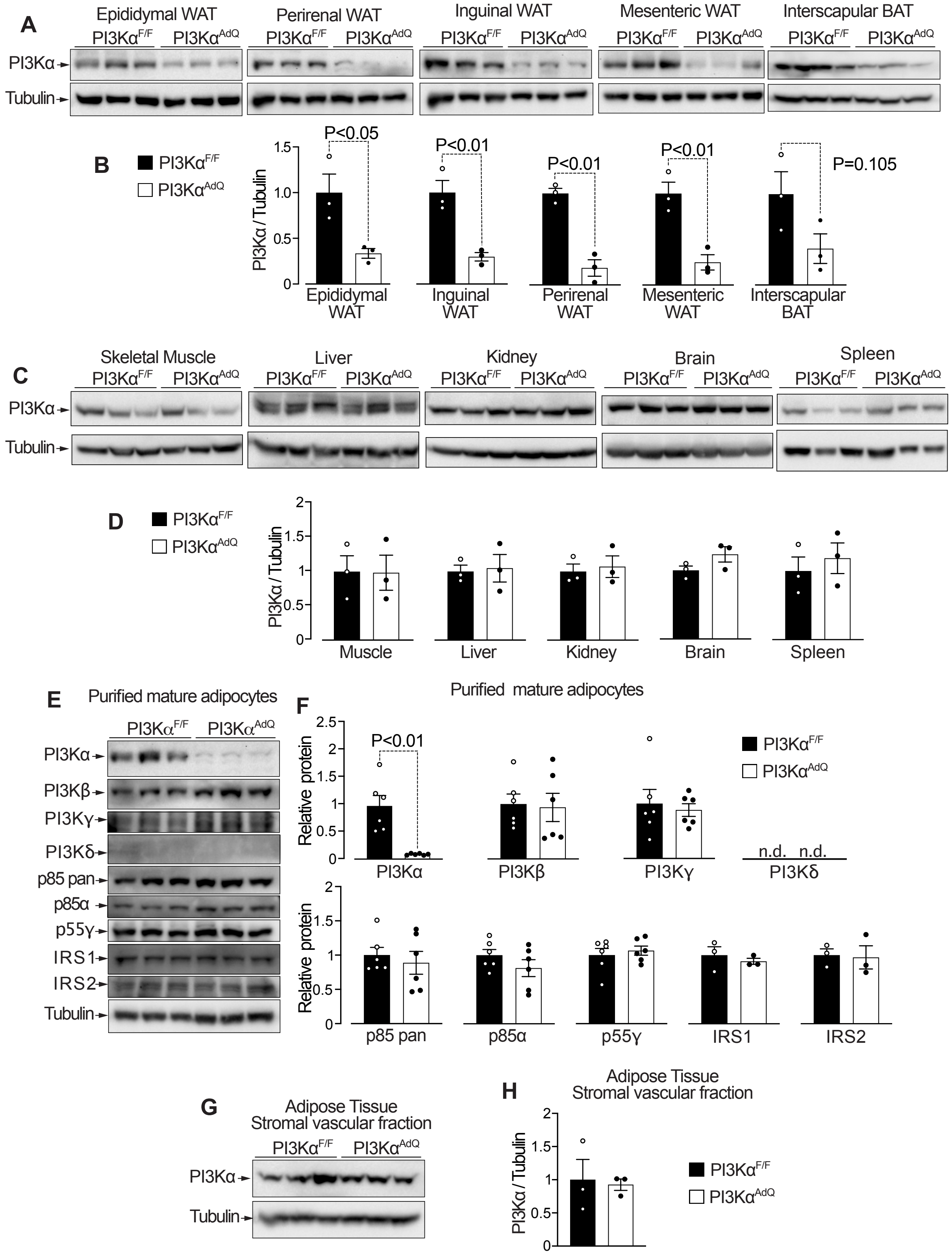
A
Epididymal WAT
Perirenal WAT
Inguinal WAT
Mesenteric WAT
Interscapular BAT
PI3Kα^F/F
PI3Kα^AdQ
PI3Kα
Tubulin
B
PI3Kα / Tubulin
P<0.05
P<0.01
P<0.01
P<0.01
P=0.105
Epididymal WAT
Inguinal WAT
Perirenal WAT
Mesenteric WAT
Interscapular BAT
C
Skeletal Muscle
Liver
Kidney
Brain
Spleen
D
Muscle
Liver
Kidney
Brain
Spleen
E
Purified mature adipocytes
PI3Kβ
PI3Kγ
PI3Kδ
p85 pan
p85α
p55γ
IRS1
IRS2
F
Purified mature adipocytes
Relative protein
n.d. n.d.
G
Adipose Tissue
Stromal vascular fraction
H

**Figure S1. Efficient and specific PI3Kα deletion in PI3Kα$^{AdQ}$ mice, related to Figure 1.**
(A) PI3Kα$^{AdQ}$ and PI3Kα$^{F/F}$ mice from Figure 1 were sacrificed, and PI3Kα protein abundance in their adipose tissue pads was measured by immunoblotting. (B) Quantification of the immunoblots in A. (C) PI3Kα protein abundance in muscle, liver, kidney, brain, and spleen was measured by immunoblotting. (D) Quantification of the immunoblots in C. (E) The abundances of class-1 PI3K catalytic and adapter subunits and of the insulin receptor substrates IRS1 and IRS2 were quantified by immunoblot in mature adipocytes purified from epididymal fat pads. (F) Quantification of the immunoblots in E. (G) Immunoblot analysis of PI3Kα protein abundance in the stromal vascular fractions of the epididymal fat pads in E. (H) Quantification of the immunoblots in G. n=3 mice per group for A-D and G-H, n=6 for E-F (n=3 for IRS1 and IRS2). Data are mean ± s.e.m. Statistical analysis was performed using the unpaired t-test or Mann-Whitney for E (n=6).

**Fig. S2**

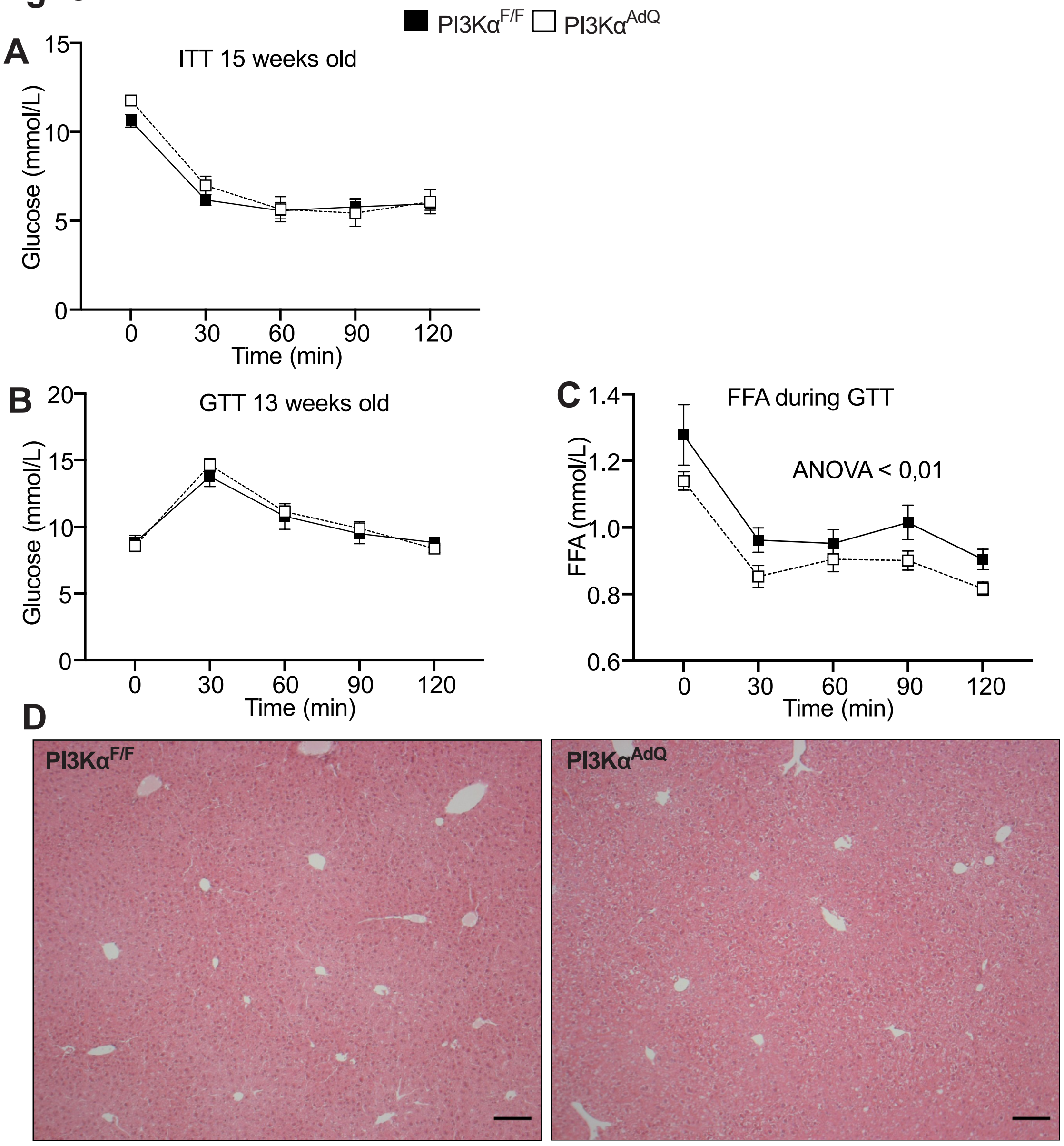


**Figure S2. PI3K$\alpha^{AdQ}$ mice display normal glucose and lipid control, related to Figure 1.** (A) Insulin tolerance test (ITT): PI3K$\alpha^{AdQ}$ and PI3K$\alpha^{F/F}$ mice were fasted for 4 hours, and blood glucose was measured at the indicated time points after i.p. injection of 1 IU/kg of insulin. (B) Glucose tolerance test (GTT): PI3K$\alpha^{AdQ}$ and PI3K$\alpha^{F/F}$ mice were fasted for 4 hours, and blood glucose was measured at the indicated time points after i.p. injection of 1 g/kg of glucose. (C) Quantification of serum-free fatty acids during the GTT. (D) Representative image of liver histology. Scale bar 100 μm. n=6 mice per group, data are mean ± s.e.m. Statistical analysis was performed using repeated measures two-way ANOVA.

Fig. S3

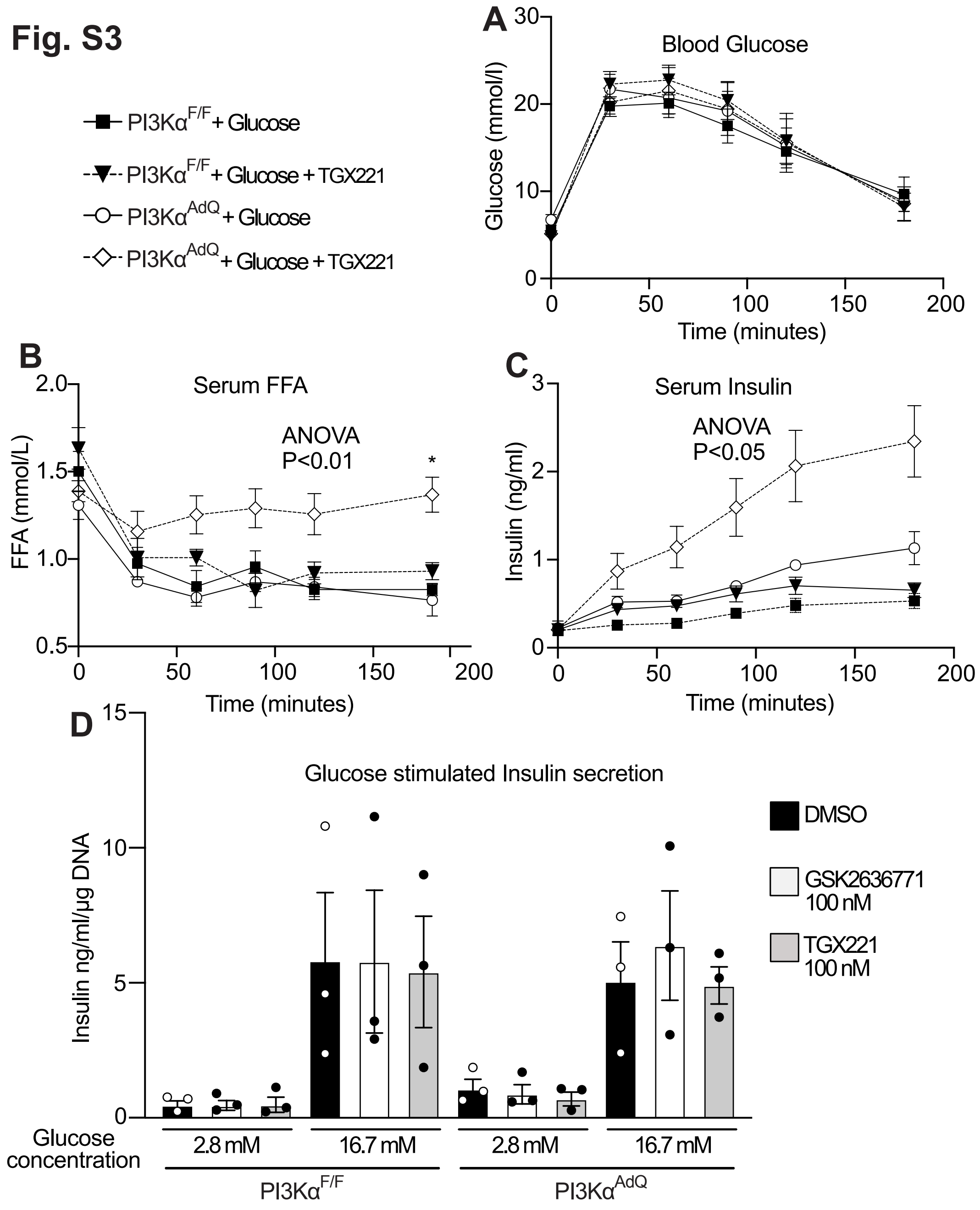


**Figure S3. Adipocyte PI3K activity links adipostasis with insulin secretion through an adipoincretin effect, related to Figure 2.**

(A) PI3Kα$^{AdQ}$ and PI3Kα$^{F/F}$ mice were fasted for 12 hours and injected i.p. with 2 g/Kg of glucose with either vehicle or 8 mg/Kg of the PI3Kβ-selective inhibitor TGX221. Blood glucose was measured at the indicated time points. (B) Serum FFA, and (C) serum insulin of the mice from

A. (D) Glucose stimulated insulin secretion of islets isolated from PI3K$\alpha^{AdQ}$ and PI3K$\alpha^{F/F}$ mice kept in the presence of either vehicle, the PI3K$\beta$-selective inhibitors GSK2636771 or TGX221. n=6 mice per group in A-C and n=3 for D. Data are mean ± s.e.m. Statistical analysis was performed using repeated measures two-way ANOVA and Šídák's multiple comparisons test for A-C, and unpaired t-test for D. For B, glucose significantly reduced FFA levels in all groups except for PI3K$\alpha^{AdQ}$ mice co-treated with TGX221. The P-value in B is for PI3K$\alpha^{AdQ}$ + Glucose + TGX221 vs PI3K$\alpha^{AdQ}$ +Glucose. C, the indicated P-value is the most conservative for the following comparisons for PI3K$\alpha^{AdQ}$ + Glucose + TGX221 vs. PI3K$\alpha^{AdQ}$ + Glucose, and PI3K$\alpha^{AdQ}$ + Glucose + TGX221 vs. PI3K$\alpha^{F/F}$ + Glucose + TGX221.

**Fig. S4**

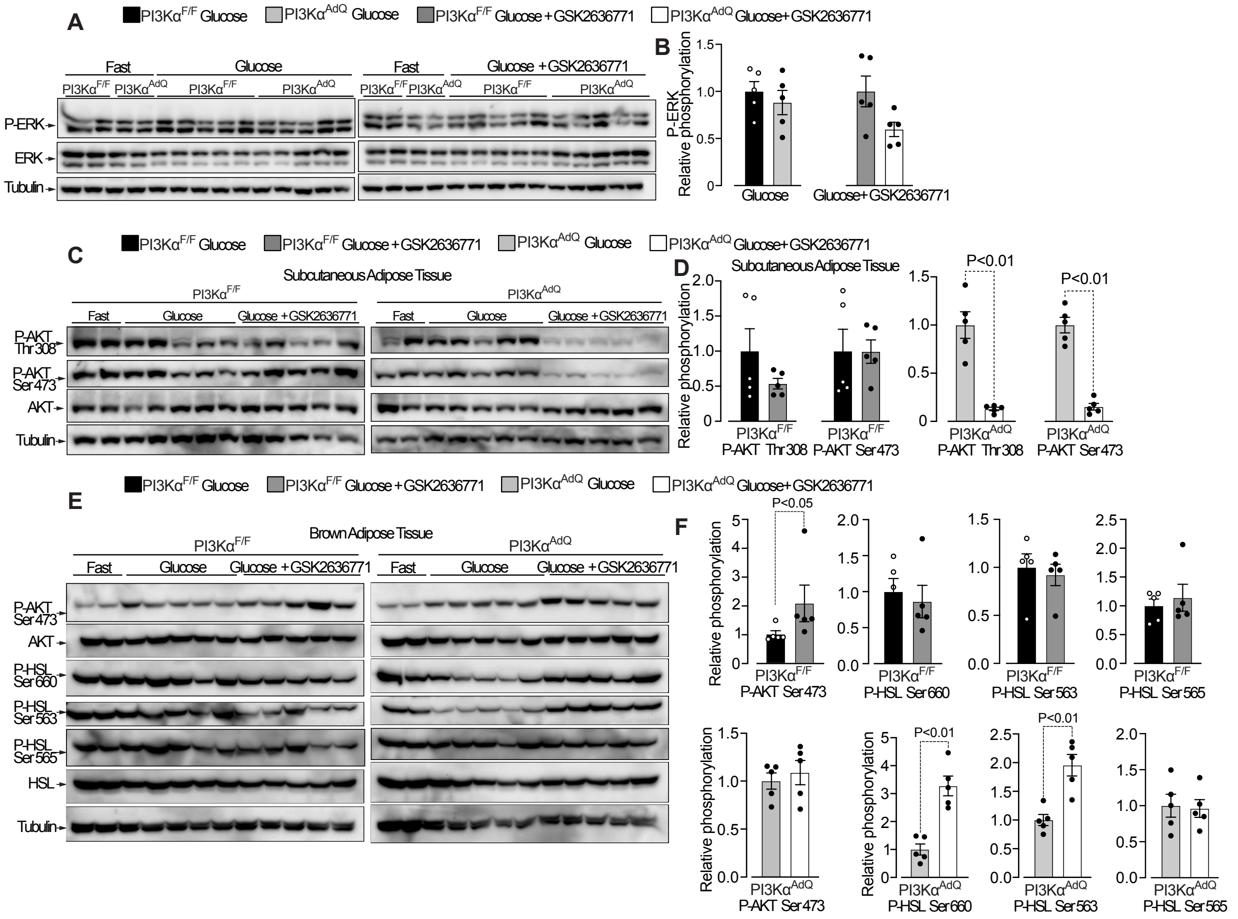

**Figure S4. Effects of adipose tissue-selective inhibition on ERK and AKT phosphorylation in fat tissues, related to Figure 2.**

(A) Immunoblot analysis of ERK phosphorylation in the epidydimal adipose tissues from PI3Kα$^{AdQ}$ and PI3Kα$^{F/F}$ mice described in Figure 2. (B) Quantification of the immunoblots in A. (C) Immunoblot analysis of AKT phosphorylation in the subcutaneous inguinal adipose tissues from PI3Kα$^{AdQ}$ and PI3Kα$^{F/F}$ mice described in Figure 2. (D) Quantification of the immunoblots in C. (E) Immunoblot analysis of AKT and HSL phosphorylation in the brown adipose tissues from PI3Kα$^{AdQ}$ and PI3Kα$^{F/F}$ mice described in Figure 2. (F) Quantification of the immunoblots in E for PI3Kα$^{F/F}$ and PI3Kα$^{AdQ}$ mice, respectively. n=2 for fasting, n=5 mice per group for all quantified datasets. Data are mean ± s.e.m. Statistical analysis was performed using the Mann–Whitney test.

**Fig. S5**

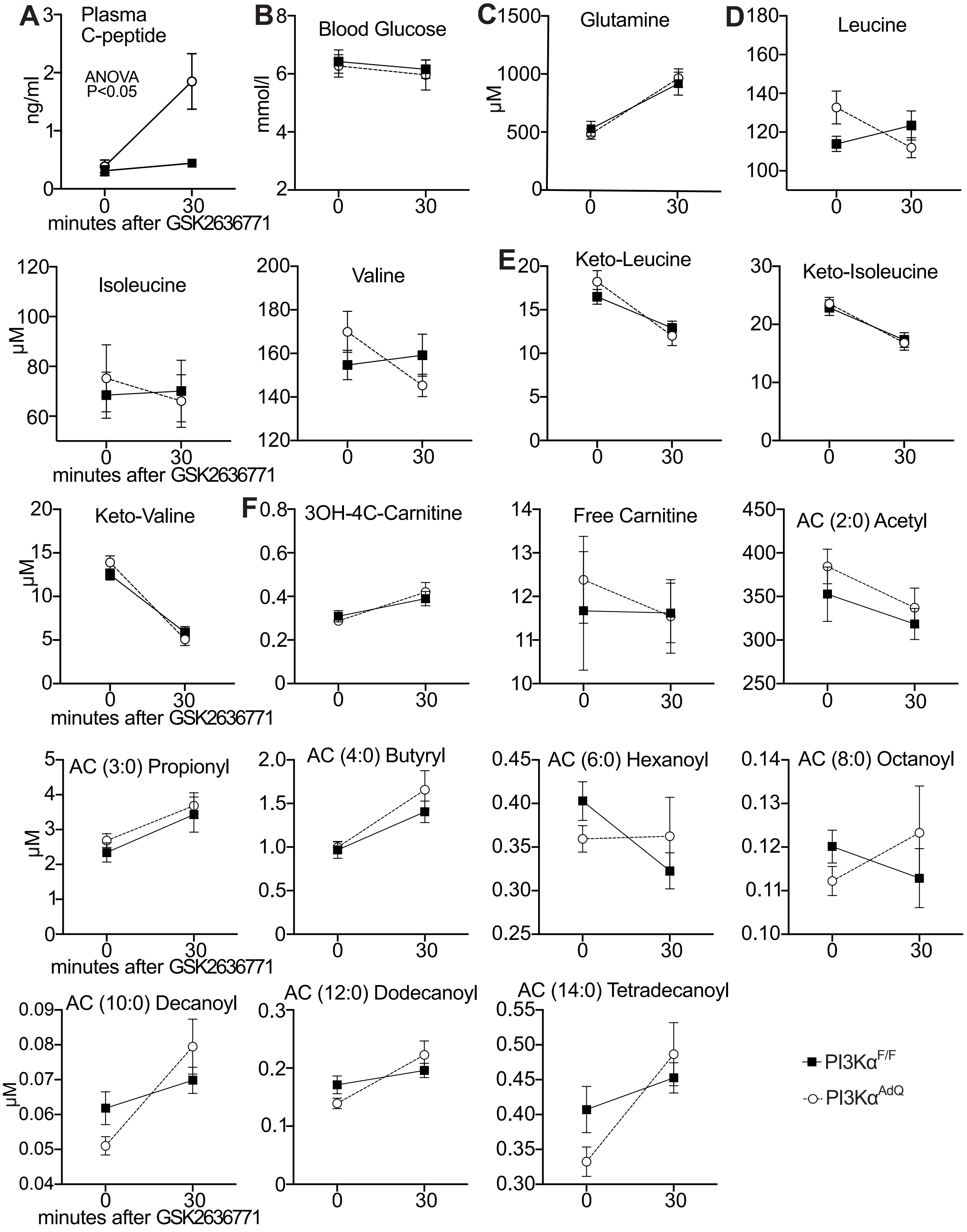

A
Plasma C-peptide
ng/ml
ANOVA P<0.05
minutes after GSK2636771
B
Blood Glucose
mmol/l
C
Glutamine
μM
D
Leucine
Isoleucine
μM
Valine
E
Keto-Leucine
Keto-Isoleucine
Keto-Valine
μM
F
3OH-4C-Carnitine
Free Carnitine
AC (2:0) Acetyl
AC (3:0) Propionyl
AC (4:0) Butyryl
AC (6:0) Hexanoyl
AC (8:0) Octanoyl
AC (10:0) Decanoyl
AC (12:0) Dodecanoyl
AC (14:0) Tetradecanoyl
PI3Kα F/F
PI3Kα AdQ

**Figure S5. Targeted metabolomics, related to Figure 5.**

PI3K$\alpha^{AdQ}$ and PI3K$\alpha^{F/F}$ mice fasted for 12 hours and were injected with GSK2636771, and plasma was collected at 0 minutes (before GSK2636771 injection) and at 30 minutes post-injection. Plasma levels of (A) C-peptide, (B) glucose, (C) glutamine, (D) branched-chain amino acids, (E) ketoacids, and (F) acylcarnitines were measured.

n=10 mice. Data are mean ± s.e.m. Statistical analysis was performed using repeated measures two-way ANOVA.

### Fig. S6

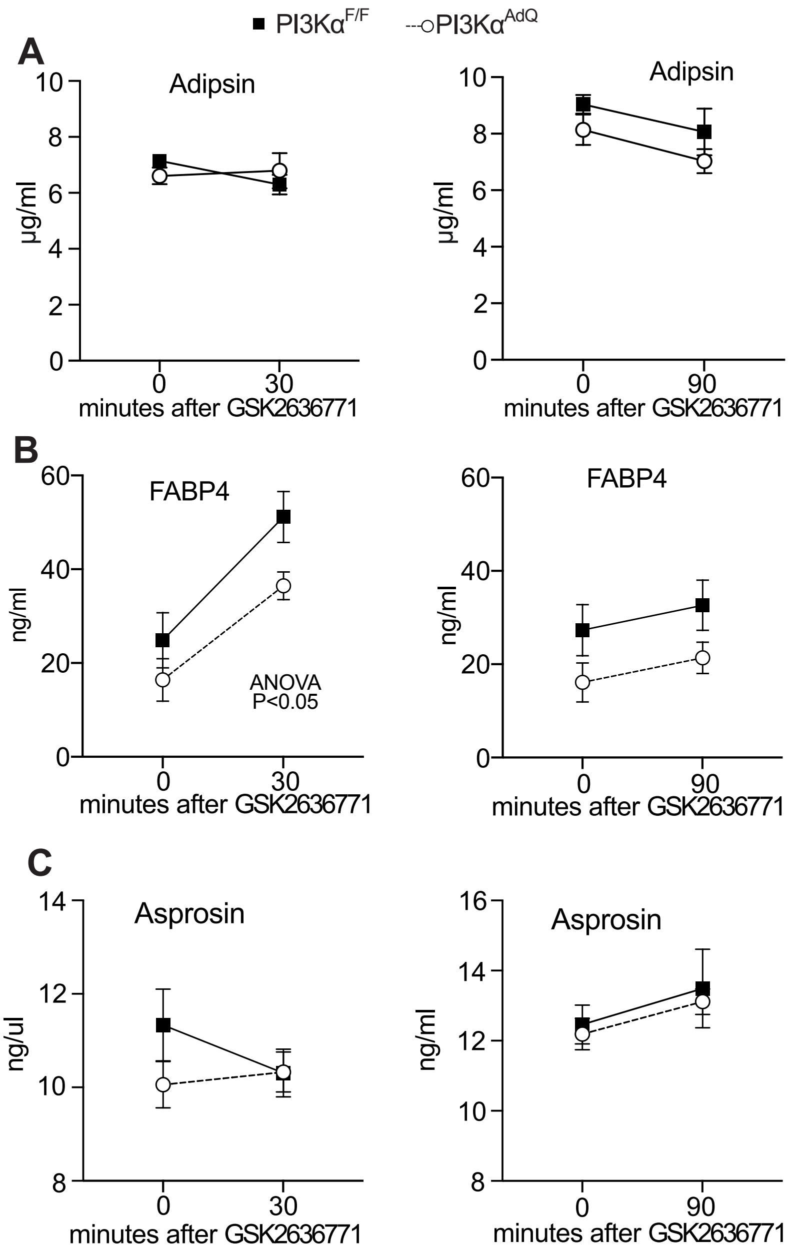


**Figure S6. The adipoincretin effect is not associated with plasma abundances of Adipsin, FABP4, or Asprosin, related to Figure 5.**

PI3Kα$^{AdQ}$ and PI3Kα$^{F/F}$ mice were fasted for 12 hours and injected with GSK2636771. Plasma was collected at 0 minutes (before GSK2636771 injection) and at 30 minutes post-injection or, in another cohort, at 0' (before GSK2636771 injection) and at 90 minutes post-injection. Plasma levels of (A) Adipsin, (B) FABP4, and (C) Asprosin were measured by ELISA.

n=7 mice for A, n=9 mice for B and n=8 for C. Data are mean ± s.e.m. Statistical analysis was performed using repeated measures two-way ANOVA. P value in B refers to an effect of GSK2636771 independent from the genotype.

Fig. S7

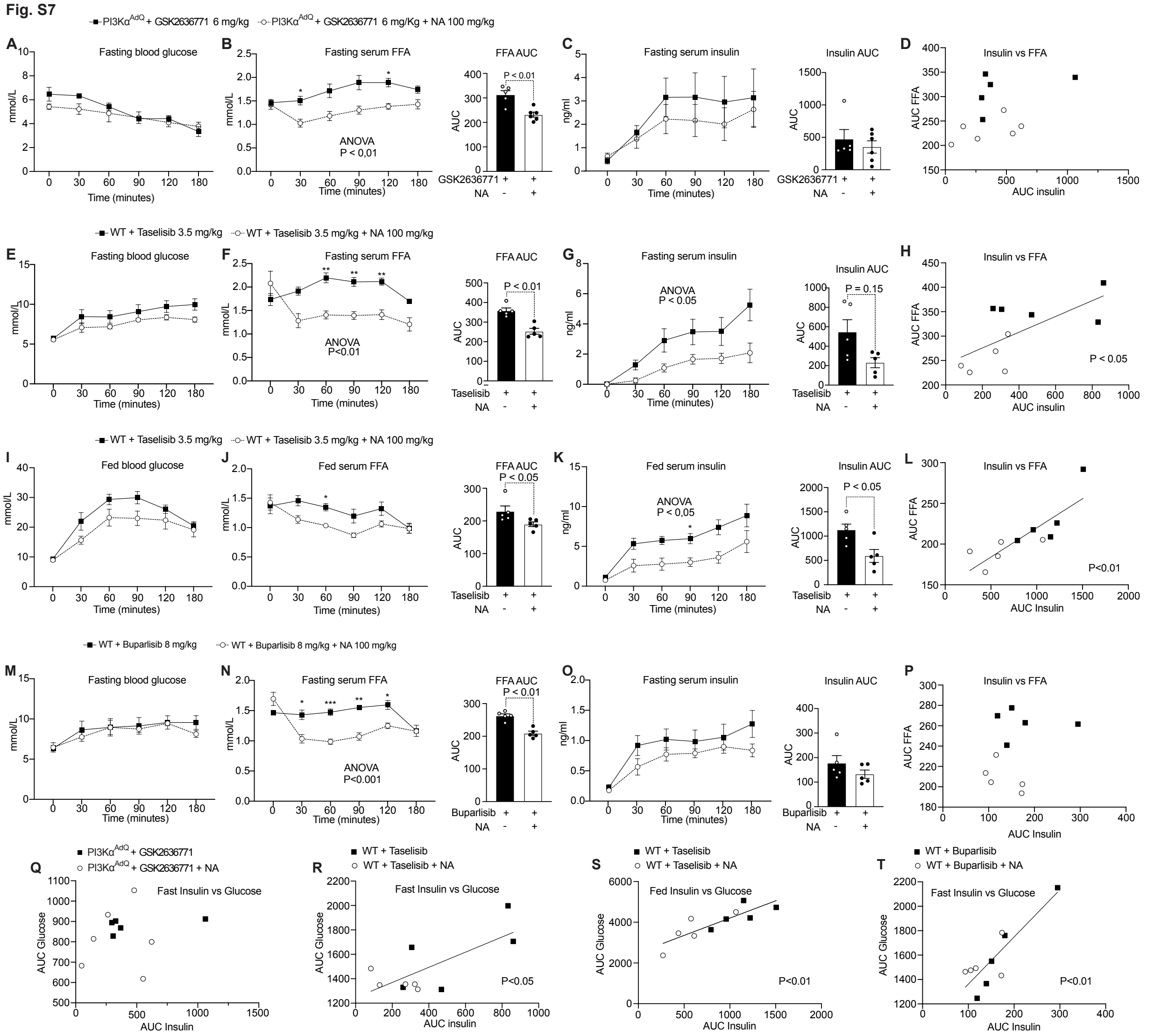


**Figure S7. Contribution of lipolysis to the adipoincretin effect and the hyperinsulinemia induced by systemic PI3K inhibition, related to Figures 6 and 7.**

PI3Kα$^{AdQ}$ mice were fasted for 12 hours and injected with GSK2636771, and one group of PI3Kα$^{AdQ}$ received GSK263771 together with nicotinic acid (NA). (A) Blood glucose, (B) serum FFA, (C) serum insulin. (D) Simple linear regression of the AUCs of insulin vs FFA from B, C. Wild-type C57BL/6 mice were fasted for 12 hours and injected with the pan-PI3K inhibitor Taselisib in the presence or absence of NA. (E) Blood glucose, (F) serum FFA, (G) serum insulin. (H) Simple linear regression of the AUCs of insulin vs FFA from F, G. The same C57BL/6 mice above were kept in fed conditions and injected with the pan-PI3K inhibitor Taselisib in the presence or absence of NA. (I) Blood glucose, (J) serum FFA, (K) serum insulin. (L) Simple linear regression of the AUCs of insulin vs FFA from J, K.

Wild-type C57BL/6 mice were fasted for 12 hours and injected with the pan-PI3K inhibitor Buparlisib in the presence or absence of NA. (M) Blood glucose, (M) serum FFA, (O) serum insulin. (P) Simple linear regression of the AUCs of insulin vs FFA from N, O.
(Q-T) Simple linear regression of the AUCs of insulin vs glucose for the mice groups in A-D, E-H, I-L, M-P, respectively.
A-D: n=5 GSK2636771 and n=6 for GSK2636771 with NA, n=5 mice for E-P.
Data are mean ± s.e.m. Statistical analysis was performed using repeated measures two-way ANOVA, Šídák's multiple comparisons test, Mann-Whitney for insulin AUC, and simple linear regression for the relationship analysis of two AUC.
